# Engineered CD47 protects T cells for enhanced antitumor immunity

**DOI:** 10.1101/2023.06.20.545790

**Authors:** Sean A. Yamada-Hunter, Johanna Theruvath, Brianna J. McIntosh, Katherine A. Freitas, Molly T. Radosevich, Amaury Leruste, Shaurya Dhingra, Naiara Martinez-Velez, Peng Xu, Alberto Delaidelli, Moksha H. Desai, Zinaida Good, Louai Labanieh, Christopher W. Mount, Yiyun Chen, Sabine Heitzeneder, Kristopher D. Marjon, Allison Banuelos, Omair Khan, Jing Huang, Savannah L. Wasserman, Jay Y. Spiegel, Sebastian Fernandez-Pol, Poul H. Sorensen, Michelle Monje, Robbie G. Majzner, Irving L. Weissman, Bita Sahaf, Elena Sotillo, Jennifer R. Cochran, Crystal L. Mackall

## Abstract

Adoptively transferred T cells and agents designed to block the CD47/SIRPα axis are promising antitumor therapeutics, which activate distinct arms of the immune system. We administered anti-CD47 (αCD47) with adoptively transferred T cells with the goal of enhancing antitumor efficacy but observed rapid macrophage-mediated clearance of T cells expressing chimeric antigen receptors (CARs) or engineered T cell receptors, which blunted therapeutic benefit. αCD47 mediated CAR T clearance was potent and rapid enough to serve as an effective safety switch. To overcome this challenge, we engineered a CD47 variant (47_E_) that engaged SIRPα and provided a “don’t-eat-me” signal that was not blocked by αCD47 antibodies. TCR or CAR T cells expressing 47_E_ were resistant to clearance by macrophages following αCD47, and mediated significant, sustained macrophage recruitment into the TME. Although many of the recruited macrophages manifested an M2-like profile, the combined therapy resulted in synergistic enhancement in antitumor efficacy. This work identifies macrophages as major regulators of T cell persistence and illustrates the fundamental challenge of combining T cell directed therapeutics with those designed to activate macrophages. It further delivers a therapeutic approach capable of simultaneously harnessing the antitumor effects of T cells and macrophages that manifests markedly enhanced potency against solid tumors.

## Introduction

Myeloid cells are the most plentiful immune cells within the tumor microenvironment (TME) and there has been great interest in therapeutically targeting them for antitumor effects^1–3^. Increased levels of tumor associated macrophages (TAMs) are linked with poorer clinical outcomes in numerous studies^4, 5^, and substantial preclinical data demonstrates that reducing or eliminating TAMs enhances responses to chemotherapy and immunotherapy^6–8^. However, dozens of clinical studies testing CSF1R and CCR2 inhibitors, designed to deplete TAMs and tumor associated myeloid cells, have been completed or are ongoing, and thus far none have demonstrated significant clinical benefit^2, 3, 9^. Alternatively, increasing TAM density is correlated with improved clinical outcomes in some cancers^5, 10^, and augmenting the phagocytic activity of TAMs by blocking the CD47/SIRPα axis mediates antitumor effects in several preclinical models^11–14^. Clinical trials of agents designed to block the CD47/SIRPα axis demonstrated antitumor activity in some liquid tumors when combined with additional agents^15, 16^, but clear evidence for single agent activity or activity in solid cancers is lacking^17, 18^. Thus, despite extensive effort, effective therapeutic approaches to target TAMs for clinical benefit remain elusive.

Adoptive T cell therapy using chimeric antigen modified T cells (CAR T) has demonstrated success in treating hematologic malignancies^19–28^, but less than 50% of patients treated with FDA approved CAR T experience durable disease control^29, 30^ and CAR T cells have been less effective in treating solid tumors^31^, which make up most cancers^32^. Resistance to adoptive T cell therapies is attributed to multiple factors, including suppressive myeloid cells within the TME^33^. Here, we sought to enhance the efficacy of adoptive T cell therapy by co-administering a blocking anti-CD47 monoclonal antibody (αCD47), based upon the hypothesis that macrophage mediated tumor phagocytosis is orthogonal to T cell mediated tumor killing and thus would provide at least an additive benefit. However, CD47 blockade completely abrogated the activity of adoptive T cell therapy through macrophage mediated depletion of the transferred T cells, in a manner sufficiently rapid and complete to serve as a safety switch in a lethal, autoreactive CAR T cell model.

We thus tested the hypothesis that selective CD47 blockade on tumor cells, but not T cells, would prevent macrophage mediated depletion of T cells and thereby enable benefit from simultaneous T cell and macrophage mediated antitumor effects. We created and expressed on T cells an engineered CD47 (47_E_) that retained SIRPα signaling but was not blocked by αCD47. Adoptive transfer of T cells expressing 47_E_ administered with αCD47 induced sustained high levels of macrophages in the TME and dramatically enhanced antitumor activity. These results demonstrate that the antagonistic effects of T cell plus macrophage targeting therapies can be converted into synergistic effects when approaches are incorporated to prevent macrophage mediated phagocytosis of T cells and provide a potential explanation for the limited therapeutic effectiveness thus far of therapies designed to either deplete TAMs or increase inflammatory macrophages within the TME.

## Results

### CD47 blockade abrogates antitumor efficacy of CAR T therapy due to T cell depletion

To test the hypothesis that augmenting macrophage phagocytosis via CD47 blockade could improve the efficacy of CAR based T cell therapy, we utilized the MG63.3 osteosarcoma model which we previously showed to be sensitive to CAR T therapy^34, 35^, and sensitive to αCD47 plus αGD2 mAb (monoclonal antibody) therapy^36^. Mice implanted orthotopically with MG63.3 cells received the αCD47 mAb B6H12^36–39^, before administering either B7H3.BBζ- or GD2.BBζ-CAR T cells (**Fig. 1a**). The results replicated the antitumor effects of each CAR T against MG63.3, but we observed no significant antitumor effect in animals receiving either CAR T therapy plus B6H12, demonstrating antagonistic activity (**Fig. 1b,c**). Similarly, while B7H3.BBζ-CAR T mediated antitumor effects against D425 orthotopic medulloblastoma xenografts^34^, we observed no antitumor effects when B7H3.BBζ-CAR T were co-administered with B6H12 (**Extended Data Fig. 1a-c**).

**Figure 1:**
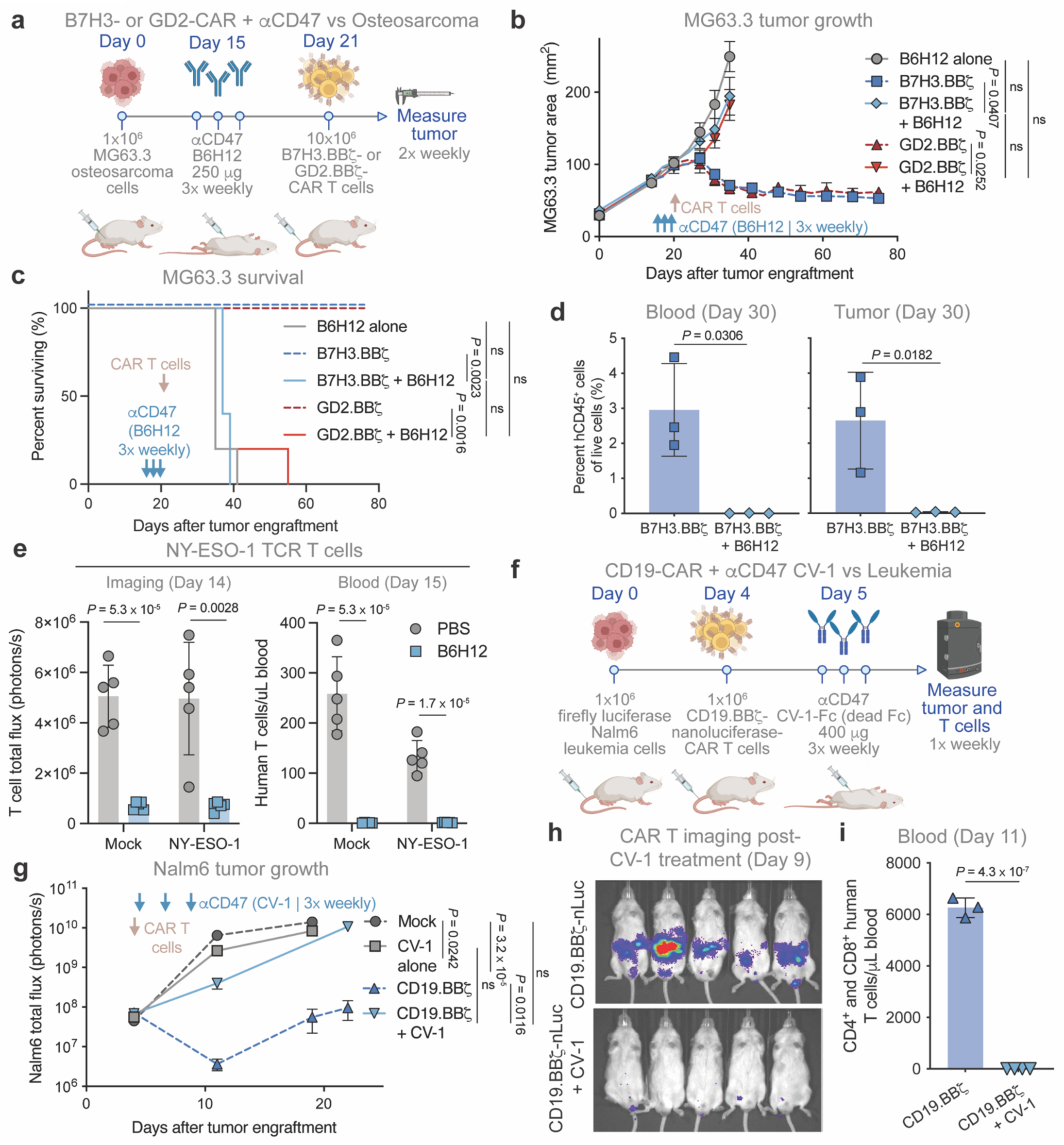
CD47 blockade leads to loss of CAR T cell efficacy *in vivo* due to T cell depletion. (a – d) MG63.3 osteosarcoma model treated with CAR T cells and B6H12. (a) MG63.3 model treatment scheme. Mice engrafted orthotopically in the tibia periosteum with 1⨉10^6^ MG63.3 were treated intraperitoneally (IP) ± B6H12 three times per week (400 µg/dose) starting on day 15, then intravenously (IV) with 10⨉10^6^ B7H3.BBζ- or GD2.BBζ-CAR T cells on day 21. (b) MG63.3 tumor growth in animals treated as described in (a). Data are the mean ± SEM of n = 5 mice/arm. (c) Survival of mice in the MG63.3 model shown in (b). n = 5 mice per treatment arm. (d) Quantification of T cells (hCD45^+^) by flow cytometry from blood and tumor in the MG63.3 model, treated IV with 10⨉10^6^ B7H3.BBζ-CAR T cells on day 15 ± 3 doses of B6H12 treatment (400 µg/dose; IP) collected on day 30 after tumor engraftment. Data are the mean ± SD of n = 3 mice. (e) Quantification of T cells by BLI (left, day 14) and in the blood by flow cytometry (hCD45^+^; right, day 15) in the A375 melanoma – NY-ESO-1 TCR model, treated as described in Extended Data Fig. 1j. Mice engrafted subcutaneously (SQ) with 3⨉10^6^ A375 were treated IV with 1⨉10^6^ mock-Antares or NY-ESO-1-Antares-TCR T cells on day 7 ± three doses of B6H12 (250 µg/dose; IP) on days 9, 11, and 14. Data are the mean ± SD of n = 5 mice. Mock is a shared control duplicated in T cell depletion in Fig. 8a & 8b. (f – i) Nalm6 leukemia model treated with CAR T cells and CV-1. (f) Nalm6 with CV-1 treatment scheme. Mice engrafted IV with 1⨉10^6^ Nalm6-fLuc were treated IV with 1⨉10^6^ CD19.BBζ-nLuc-CAR T cells on day 4, ± CV-1 (400 µg/dose; IP) three times per week starting on day 5. (g) Nalm6 – CV-1 model tumor growth, treated as described in (f). Data are the mean ± SEM of n = 5 mice/arm. (h) T cell BLI on day 9 of mice treated as described in (f). (i) T cells in the blood (hCD4^+^ and hCD8^+^) by flow cytometry on day 11 of mice treated as described in (f). Data are the mean ± SD of n = 3 mice for the CD19.BBζ group and n = 4 for the CD19.BBζ + CV-1 group. [(b), (g)] Two-way analysis of variance (ANOVA) test with Tukey’s multiple comparison test. ns = not significant. [(c)] Log-rank Mantel-Cox test. ns = not significant. [(d), (e), and (i)] Unpaired two-tailed Student’s *t* test. See also Extended Data Figures 1 and 2.

To interrogate the cause of therapeutic failure with dual treatment, we analyzed T cell levels in mice bearing MG63.3 tumors 15 days after B7H3.BBζ-CAR T ± B6H12. Strikingly, while human T cells were detectable in tumor and blood in CAR only treated mice, they were absent in mice co-treated with B6H12 (**Fig. 1d**, **Extended Data Fig. 1d**). To characterize the kinetics of CAR T cell depletion *in vivo* in animals treated with adoptive T cell transfer plus B6H12, we monitored levels of CD19.28ζ-CAR T expressing nanoluciferase (CD19.28ζ-nLuc) via bioluminescent imaging (BLI) in animals inoculated with Nalm6 leukemia incorporating firefly luciferase (Nalm6-fLuc) ± B6H12 (**Extended Data Fig. 1e**)^35^. Consistent with the results in the MG63.3 and D425 models, B6H12 completely ablated CD19.28ζ-CAR T efficacy (**Extended Data Fig. 1f,g**) and BLI demonstrated a significant loss of CAR T signal shortly after B6H12 treatment (**Extended Data Fig. 1h**). CAR T cells in the dual treatment group remained undetectable via BLI over the course of three weeks and were absent from spleens of mice at the conclusion of the experiment (**Extended Data Fig. 1h,i**). Of note, the modest single agent efficacy of B6H12 in this system was not impacted by co-administration of CD19.28ζ-CAR T cells (**Extended Data Fig. 1f**).

We next interrogated pairing of αCD47 with adoptive transfer of T cells engineered to express the widely validated NY-ESO-1 targeting TCR^40^. We implanted mice with A375 melanoma cells, which display the NY-ESO-1 peptide, in a flank xenograft model and administered mock T cells, or NY-ESO-1-TCR T cells, ± B6H12 (**Extended Data Fig. 1j**). All T cells also expressed Antares, a nanoluciferase-orange fluorescent protein fusion with enhanced sensitivity for BLI^35, 41^. As observed for CAR T cells, mock and NY-ESO-1-TCR T cells were depleted to undetectable levels after B6H12 administration (**Fig. 1e**, **Extended Data Fig. 1k**), despite robust levels pre-treatment (**Extended Data Fig. 1l**). Together, the data demonstrate that αCD47 induces rapid and near-complete depletion of adoptively transferred T cells that are not genetically modified, cells engineered to express a transgenic TCR and cells engineered to express CARs spanning differing targeting domains (B7H3, GD2, and CD19) and costimulatory domains (4-1BB and CD28).

### CD47 provides an essential “don’t eat me” signal on CAR T cells to prevent macrophage mediated phagocytosis

We next sought to determine whether T cell ablation in these models was due to antibody dependent phagocytosis that requires FcR engagement^42, 43^ by administering CV-1^44^, an engineered SIRPα Fc-fusion, which contained an immunologically inert hIgG1 Fc-domain with LALA-PG mutations^43^. Similar to results with B6H12, CV-1 co-treatment with CD19.BBζ-CAR T led to complete loss of CAR T efficacy in Nalm6-fLuc bearing mice (**Fig. 1f,g, Extended Data Fig. 2a,b**), and near total T cell depletion (**Fig. 1h,i, Extended Data Fig. 2c**). To further assess the requirement of CD47 expression for survival of adoptively transferred T cells, we generated CD47 knock-out (47_KO_) T cells via CRISPR/Cas9 and confirmed loss of αCD47 and soluble SIRPα binding (**Fig. 2a**), which was restored upon exogenous expression of full-length, wild-type CD47 in 47_KO_ cells (47_WT_). We next treated Nalm6-fLuc bearing mice with 47_KO_- or 47_WT_-CD19.28ζ-nLuc-CAR T and observed that 47_WT_-CAR T cells expanded *in vivo*, mediated robust tumor control and significantly prolonged survival, whereas 47_KO_-CAR T were depleted and delivered no anti-tumor activity or benefit in survival (**Fig. 2b-d, Extended Data Fig. 2d**). Even in the absence of tumor, we observed robust expansion of 47_WT_-CAR T, while 47_KO_-CAR T were depleted to a similar degree as 47_WT_-CAR T after B6H12 administration (**Extended Data Fig. 2e-i)**.

**Figure 2:**
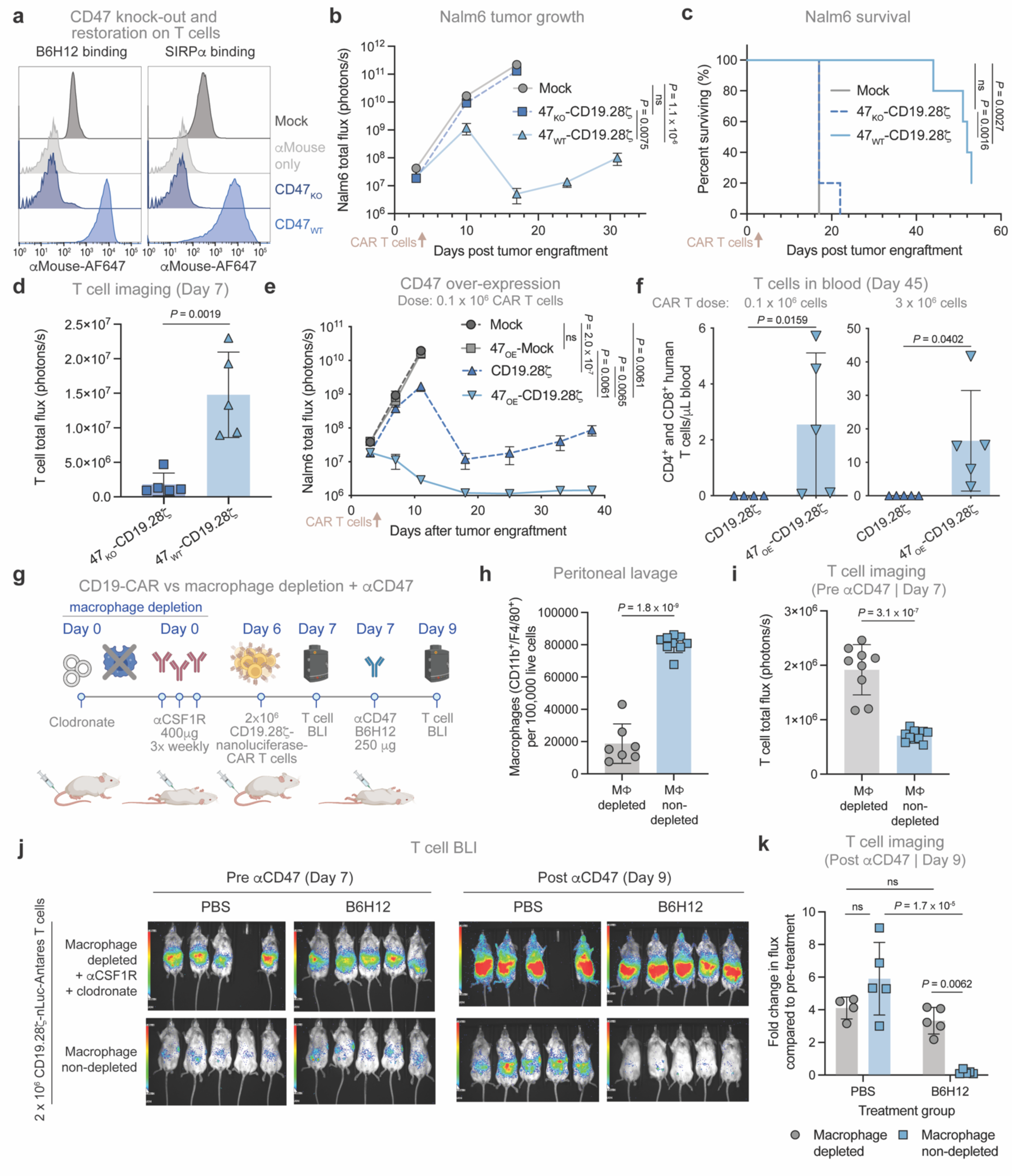
CD47 expression on CAR T cells prevents macrophage mediated depletion via Fc independent phagocytosis. (a) CD47 knock-out (CD47_KO_) efficiency in primary human T cells by flow cytometry. CD47_WT_ cells are CD47_KO_ with exogenous expression of wild-type CD47. Data are representative of n > 3 donors. (b – d) Nalm6 leukemia model treated with 47_KO_-CAR T cells. (b) Quantification of Nalm6 tumor growth by BLI. Mice engrafted IV with 1⨉10^6^ Nalm6-fLuc were treated IV with 0.15⨉10^6^ mock, 47_KO_- or 47_WT_-CD19.28ζ-nLuc-CAR T cells on day 4. 47_WT_ cells are 47_KO_ with exogenous expression of WT CD47. Data are the mean ± SEM of n = 5 mice/arm. (c) Survival of Nalm6 bearing mice shown in (b). n = 5 mice per treatment arm. (d) Quantification of T cell BLI on day 7 post CAR T as described in (b). Data are the mean ± SD of n = 5 mice. (e – f) Overexpression of CD47 (47_OE_) on CAR T cells. (e) Quantification of Nalm6 tumor growth by BLI. Mice engrafted IV with 1⨉10^6^ Nalm6 were treated IV with 0.1⨉10^6^ mock, 47_OE_-mock, CD19.28ζ-, or 47_OE_-CD19.28ζ-CAR T cells on day 4. Data are the mean ± SEM of n = 5 mice/arm. Mock and 47_OE_-mock are shared conditions duplicated in Extended Data Fig. 2l. (f) Quantification of CD4^+^ and CD8^+^ T cells on day 45 after CAR T administration in the blood of mice treated with 0.1⨉10^6^ (left) or 3⨉10^6^ (right) CD19.28ζ- or 47_OE_-CD19.28ζ-CAR T cells, by flow cytometry. Data are the mean ± SD of n = 5 mice. (g – k) Macrophage depletion model. (g) Macrophage depletion scheme. Non-tumor bearing mice were treated with clodronate (200 µL; IV) on day 0 and αCSF1R (400 µg/dose; IP) three times per week starting on day 0 to deplete macrophages. Mice were then treated IV with 2⨉10^6^ CD19.28ζ-nLuc-CAR T cells on day 6, followed by a single 250 µg IP dose of B6H12 on day 7. Mice were imaged by BLI before (day 7) and after (day 9) αCD47 treatment. (h) Quantification of CD11b^+^/F4/80^+^ macrophages collected via peritoneal lavage by flow cytometry on day 13. Data are the mean ± SD of n = 7 (macrophage depleted) or n = 9 (macrophage non-depleted) mice. (i) Quantification of T cell BLI one day after IV administration of 2⨉10^6^ CD19.28ζ-nLuc-CAR T cells (day 7). Data are the mean ± SD of n = 9 (macrophage depleted) or n = 10 (macrophage non-depleted) mice. (j) T cell BLI before and after B6H12 treatment of mice treated as described in (g) (k) Fold change in quantified T cell BLI after treatment with B6H12 of mice treated as descried in (g). Data are the mean ± SD of n = 4 (macrophage depleted, PBS treated) or n = 5 (all other groups) mice. [(b), (e), and (k)] Two-way ANOVA test with Tukey’s multiple comparison test. ns = not significant. [(c)] Log-rank Mantel-Cox test. [(d), (f; right panel), (h), and (i)] Unpaired two-tailed Student’s *t* test. [(f; left panel)] Mann-Whitney test. See also Extended Data Figure 2.

Given the critical importance of CD47 expression for T cell persistence *in vivo*, we wondered whether overexpression of CD47 (47_OE_), a technique that has been reported to prevent immune rejection by allogeneic cells^45^, could enhance CAR T cell persistence and efficacy in a model where immune rejection does not occur due to the profound immune suppression of NSG mice. We treated Nalm6-fLuc bearing mice with a subtherapeutic dose of CD19.28ζ-CAR T or mock cells ± 47_OE_ (**Extended Data Fig. 2j**). 47_OE_-CD19.28ζ-CAR T mediated significantly better long-term antitumor efficacy compared to control CAR T cells (**Fig. 2e**, **Extended Data Fig. 2k,l**), and manifested improved persistence 45 days after administration (**Fig. 2f**). Thus, CD47 overexpression enhances CAR T persistence and efficacy, even in settings where immune rejection does not occur. Together these data demonstrate that survival of adoptively transferred T cells require CD47 expression and SIRPα engagement and that CD47 expression density by T cells substantially impacts survival and antitumor potency of T cells, independent of the previously reported effects of CD47 on allograft rejection.

Based upon the evidence that CD47 blockade enhances macrophage phagocytosis of tumors^14, 46, 47^, we queried whether macrophage mediated T cell depletion induced by αCD47 treatment was responsible for the findings described above. Mice were treated with clodronate plus αCSF1R mAb to deplete macrophages^36^ (**Fig. 2g,h**, **Extended Data Fig. 2m**), then received CD19.28ζ-nLuc-CAR T cells ± B6H12 (**Fig. 2g**). On Day 7, one day following adoptive transfer but prior to B6H12 administration, we observed significantly higher T cell BLI in mice with depleted macrophages consistent with macrophage mediated depletion of T cells even in the absence of αCD47 (**Fig. 2i,j**). Upon B6H12 administration, we observed no loss of CAR T signal by BLI in mice with depleted macrophages, compared to a significant reduction in T cell BLI in mice with an intact macrophage compartment (**Fig. 2j,k**). Together, these results identify macrophages as barriers to engraftment and antitumor efficacy of adoptively transferred T cells, and demonstrate an essential requirement for adequate levels of CD47 on T cells to engage SIRPα, even in hosts incapable of recognizing allogeneic disparities. They further explain the futility of combining αCD47 with adoptive T cell therapy and implicate macrophage mediated phagocytosis as an important regulator of T cell persistence *in vivo*.

### Human CAR T cells are robustly phagocytosed by human macrophages in vitro, and clinical data provides evidence of macrophage mediated phagocytosis of CAR T cells

We next investigated the potential for primary human macrophages to phagocytose primary human T cells *in vitro*. We observed low levels of macrophage phagocytosis of mock transduced T cells at baseline, while T cells transduced to express CARs were phagocytosed at significantly higher levels (**Fig. 3a**), which was further increased after addition of B6H12 (**Fig. 3a**). Phagocytosis by macrophages is regulated by a balance of “eat me” signals, such as calreticulin^47^, and “don’t eat me” signals, such as CD47. Using flow cytometry, we observed that CAR T cell expression of calreticulin increased over time in culture (**Fig. 3b,c**), while CD47 expression decreased during the same period (**Fig. 3d**), consistent with a model wherein aged CAR T cells are more susceptible to phagocytosis.

**Figure 3:**
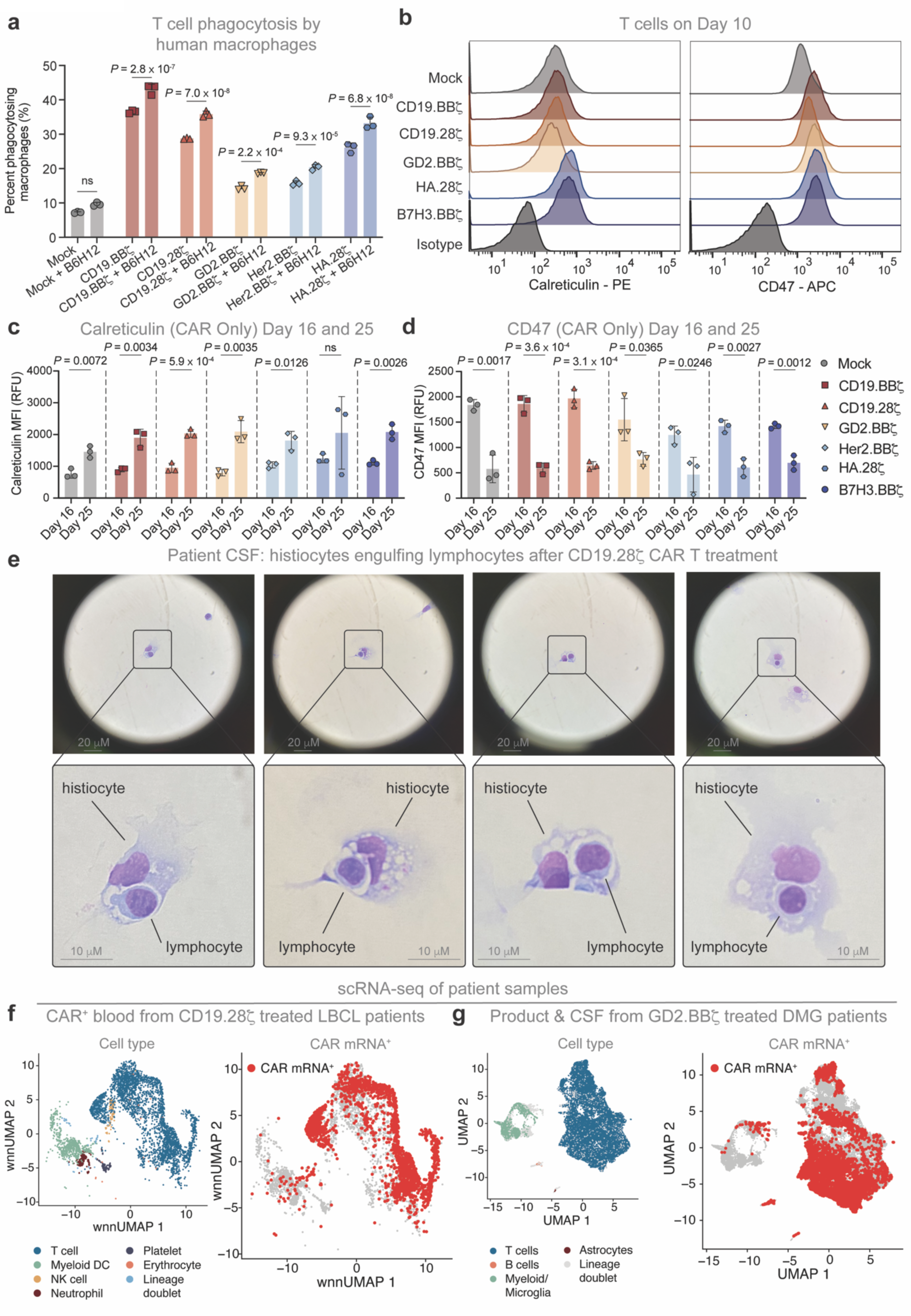
CAR T cells are phagocytosed by macrophages *in vitro* and in patients. (a) Quantification of phagocytosis of CSFE labeled CAR T cells by primary human macrophages by flow cytometry, following one hour of co-culture. Data are the mean ± SD of n = 3 triplicate wells. Data is reproducible across four different macrophage donors. (b) Calreticulin (left) and CD47 (right) expression on CAR T cells by flow cytometry on day 10 of culture. Data are representative of n = 3 donors. (c and d) Quantification of (c) calreticulin and (d) CD47 expression on CAR T cells on day 16 and day 25 of culture by flow cytometry. Data are the mean ± SD of n = 3 donors. (e) Microscope images of histiocytes engulfing lymphocytes collected from the CSF of a CD19.28ζ-CAR T treated large B cell lymphoma (LBCL) patient and stained with hematoxylin and eosin (1000x magnification). Lower panels are enlargements of the boxed regions of the respective upper panels. (f) Weighted nearest neighbor (wnn) UMAP single cell landscapes color coded for cell type (left) or CAR mRNA (right). Data are derived from *n* = 6,316 CAR^+^ cells sorted from the blood collected on day 7 after CAR T infusion of *n* = 9 axicabtagene ciloleucel (axi-cel; CD19.28ζ) treated LBCL patients, sampled to 500 cells/patient sample. (g) UMAP single cell landscapes color coded for cell type (left) or CAR mRNA (right). Data are derived from *n* = 65,598 cells from the manufacturing products (*n* = 40,000 cells) and CSF (*n* = 25,598 cells) of *n* = 4 GD2.BBζ treated diffuse midline glioma (DMG) patients, sampled to 500 cells/patient sample. [(a), (c), and (d)] Unpaired two-tailed Student’s *t* test. ns = not significant.

During the course of these experiments, routine cytologic analysis of cerebrospinal fluid collected from a patient treated with axicabtagene ciloleucel (axi-cel), a standard of care CD19.28ζ-CAR T cell therapy^29^, revealed histiocytes engulfing lymphocytes (**Fig. 3e**), consistent with macrophage mediated phagocytosis. To address this possibility more systematically, we analyzed single-cell RNA sequencing (scRNA-seq) data collected from two recent clinical studies that enrolled patients treated with axi-cel^48^ and GD2.BBζ-CAR T^49^, respectively. Both datasets provided clear evidence for CAR mRNA in myeloid cells, consistent with macrophage mediated phagocytosis of CAR T cells in humans (**Fig. 3f,g**). These data provide further evidence in support of a model wherein myeloid cells phagocytose CAR T cells, and thus may limit durable engraftment of adoptively transferred cells or survival of activated T cells in clinical settings.

### CD47 blockade can be utilized as a safety switch to control CAR T mediated toxicity

Because CD47 blockade led to rapid and robust CAR T cell depletion *in vivo*, we reasoned that treatment with αCD47 could be harnessed as an off-the-shelf safety switch to mitigate CAR T toxicity. To test this, we developed a toxicity model based on an integrin targeting CAR derived from an engineered cystine-knot peptide (knottin) called polyspecific integrin-binding peptide (PIP)^50^ that binds tumor-associated integrins expressed on a wide array of malignancies^51–53^ (**Fig. 4a**). We hypothesized that PIP’s broad range of selective tumor binding could result in a promising CAR design. We saw potent antitumor activity from PIP-CAR T *in vitro* (**Fig. 4b, Extended Data Fig. 3a**), but PIP-CAR T proved toxic *in vivo* (**Fig. 4c**), leading to cytokine release (**Fig. 4d**) and rapid weight loss (**Extended Data Fig. 3b**) due to on-target, off-tumor toxicity in the lungs of treated mice (**Extended Data Fig. 3c-g**). We thus treated mice with PIP.28ζ-nLuc-CAR T ± three doses of B6H12 over three successive days starting two days later (**Fig. 4e**). PIP.28ζ-CAR T treated mice quickly lost weight and demonstrated acute toxicity, with rapid T cell expansion by BLI and elevated human cytokines in the blood (**Fig. 4f-j**), however PIP.28ζ-CAR T recipients treated with B6H12 did not demonstrate any overt toxicity or weight loss, including no detectable human cytokines in the blood, due to T cell depletion (**Fig. 4f-j**). Thus, CD47 blockade shows promise as an off-the-shelf safety switch to quickly eliminate CAR T associated toxicity, as evidenced by rescue in a stringent toxicity model.

**Figure 4:**
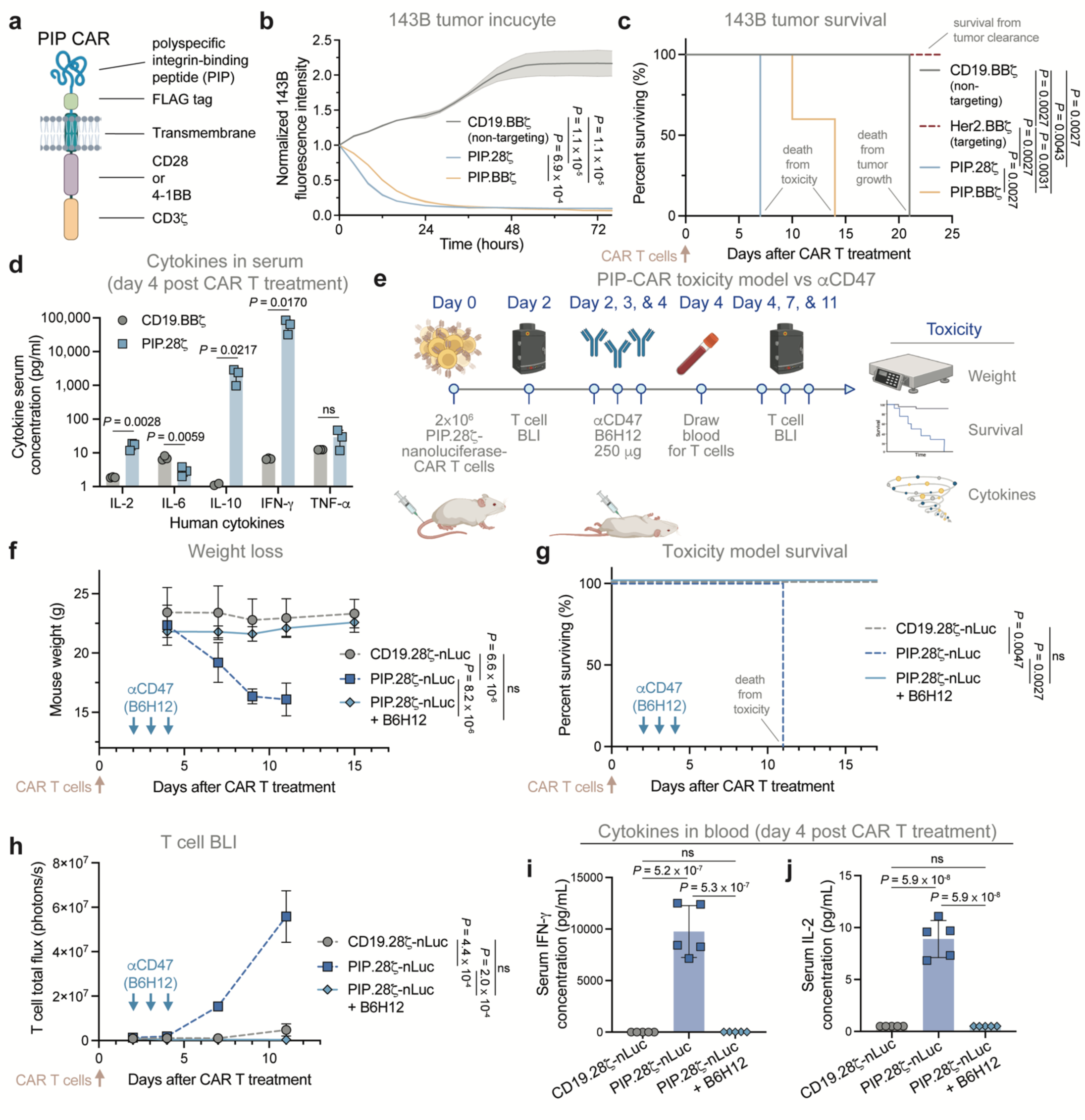
αCD47 therapy can be used as a safety switch. (a – d) PIP-CAR toxicity model (a) Cartoon of the PIP-CAR design. (b) 143B-GFP tumor killing by PIP.28ζ-, PIP.BBζ-, or non-tumor targeting CD19.BBζ-CAR T cells at a 1:1 E:T ratio measured via Incucyte assay. Data are mean ± SD of n = 3 triplicate wells and are reproducible in three independent experiments with different donors. (c) 143B osteosarcoma model survival. Mice engrafted orthotopically in the tibia periosteum with 0.5⨉10^6^ 143B were treated IV with 10⨉10^6^ CD19.BBζ-(non-tumor targeting control), Her2.BBζ-(tumor targeting control), PIP.28ζ-, or PIP.BBζ-CAR T cells on day 5. n = 5 mice per treatment arm. (d) Quantification of human cytokines in the blood of 143B bearing mice on day 4 after treatment with 10⨉10^6^ CD19.BBζ-(non-tumor targeting) or PIP.28ζ-CAR T by ELISA. Data are the mean ± SD of n = 3 mice. (e – j) PIP-CAR toxicity model treated with αCD47 (e) PIP-CAR toxicity model scheme. Mice were treated IV with 2⨉10^6^ CD19.28ζ-nLuc- or PIP.28ζ-nLuc-CAR T cells ± B6H12 treatment (250 µg daily; IP) three times, two days later. Mice were imaged by BLI before (day 2) and after (days 4, 7, and 11) B6H12 treatment. Blood was collected on day 4 for cytokine analysis. (f) Mouse weights following CAR T treatment ± B6H12, treated as described in (e). Data are mean ± SD of n = 5 mice/arm. (g) Survival of mice in the PIP-CAR toxicity model treated as described in (e). n = 5 mice per treatment arm. Data are representative of two independent experiments. (h) Quantification of T cell BLI following CAR T treatment ± B6H12, treated as described in (e). Data are mean ± SEM of n = 5 mice/arm. (i and j) Quantification of (i) IFNɣ and (j) IL-2 in the blood collected on day 4 of CAR T ± B6H12 mice, treated as described in (e) via ELISA. Data are mean ± SD of n = 5 mice. [(b), (f), and (h)] Two-way ANOVA test. ns = not significant. [(c), (g)] Log-rank Mantel-Cox test. ns = not significant. [(d)] Unpaired two-tailed Student’s *t* test. ns = not significant. [(i) and (j)] One-way ANOVA test with Tukey’s multiple comparison test. ns = not significant. See also Extended Data Figure 3.

Prolonged CAR T persistence can lead to graft versus host disease (GvHD) in xenograft models^54–56^. To determine the longevity of CAR T cell depletion following αCD47 administration, we monitored Nalm6 bearing mice treated with CD19.BBζ-CAR T ± three doses of B6H12 over three successive days for development of GvHD after successful tumor clearance (**Extended Data Fig. 3h**). After forty-eight days, we observed onset of GvHD in CD19.BBζ-CAR T mice, evidenced by alopecia and weight loss^54^ (**Extended Data Fig. 3i**). However, mice treated with CAR T plus B6H12 did not develop GvHD over the same time frame, remaining healthy with smaller spleens, and fewer lymphocyte infiltrates into the skin than CAR T only treated mice (**Extended Data Fig. 3j,k**). Thus, after short-term administration of αCD47, CAR T cells remain depleted long term and do not recur, even in situations where CAR T cells traditionally cause GvHD.

### An engineered variant of CD47 (47_E_) with selective binding retains “don’t eat me” function but does not bind αCD47 mAbs

To induce selective tumor phagocytosis via CD47 blockade while protecting T cells from phagocytosis, we sought to engineer a CD47 variant with mutations that ablate αCD47 binding while retaining binding to SIRPα (**Fig. 5a**). We first displayed the CD47 Ig-like domain on the surface of yeast and detected strong binding to B6H12, but not SIRPα (**Extended Data Fig. 4a-d**), as had been seen previously due to the lack of a free N-terminus on CD47 when displayed on yeast^57^. As a proxy for SIRPα binding, we thus used the engineered SIRPα variant, CV-1^44^ (**Extended Data Fig. 4e**) and subjected a library of yeast displayed CD47 mutant variants to six total successive FACS sorts, alternating between negative sorts against B6H12 and positive sorts towards CV-1 (**Fig. 5b**, **Extended Data Fig. 4f**). The bulk library population collected after the final sort retained binding to CV-1 but demonstrated near complete loss of binding to B6H12 (**Extended Data Fig. 4g**), and sequencing revealed that all CD47 variants contained a single A30P or Q31P point mutation (**Fig. 5c**), both of which localize to the BC loop of CD47. When displayed as individual CD47 variants on yeast, we confirmed that both A30P (47_A30P_) and Q31P (47_Q31P_) mutations demonstrated no binding to 1 µM B6H12 but manifested similar or even enhanced binding to CV-1 (**Fig. 5d**).

**Figure 5:**
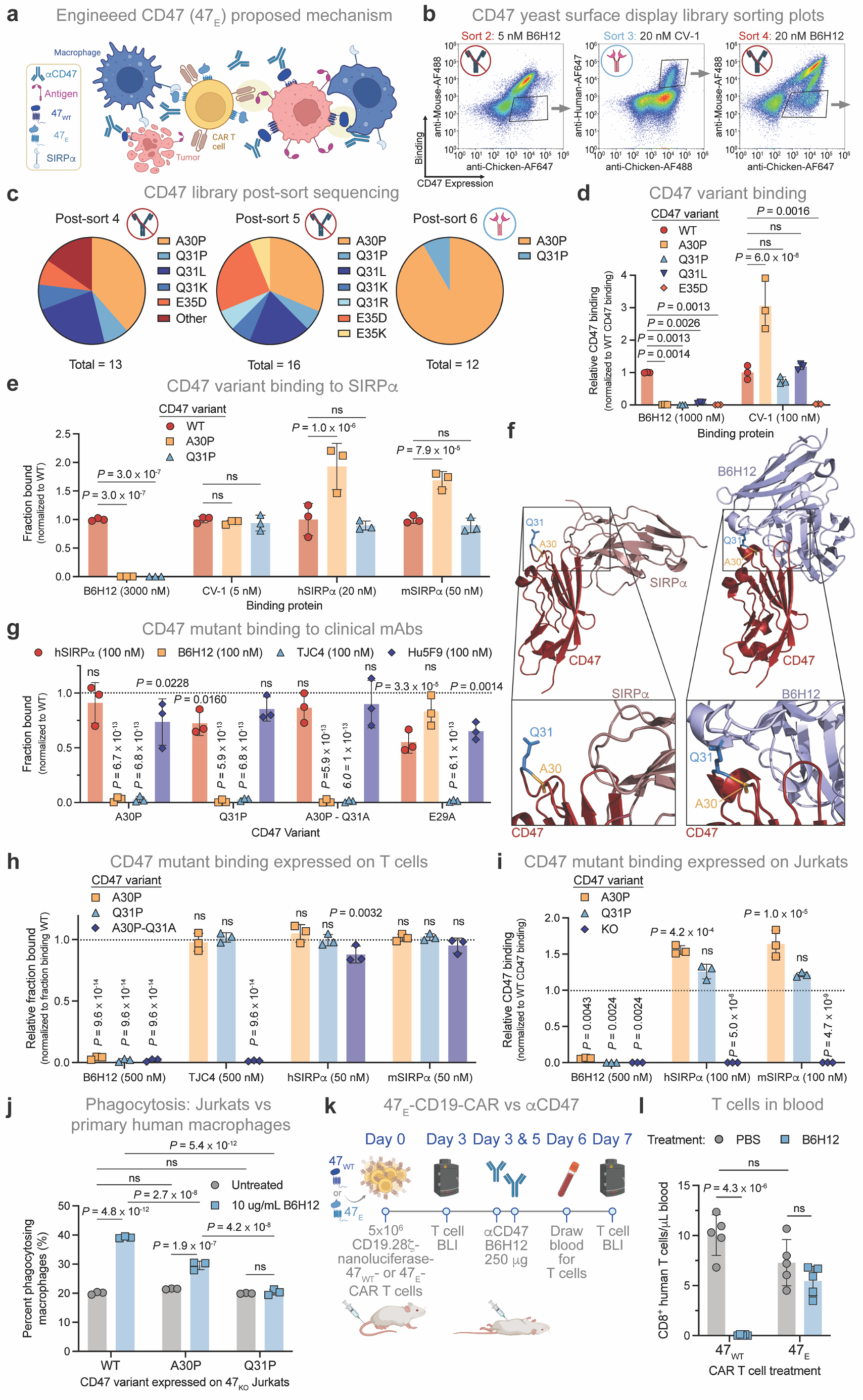
An engineered variant of CD47 retains binding to SIRPα, but no longer binds αCD47 antibodies. (a) Schematic of the engineered CD47 (47_E_) mechanism, whereby αCD47 antibodies bind tumor cells, but not 47_E_-T cells, triggering tumor-specific phagocytosis. (b) Representative positive and negative sort plots of the CD47 yeast surface display library. Black boxes indicate collected populations. (c) Consensus mutations identified in yeast sequenced after sorts 4, 5, and 6. Data are frequencies of identified mutations out of n = 13, n = 16, and n = 12 sequenced clones for sorts 4, 5, and 6, respectively. (d) Binding of B6H12 and CV-1 to CD47 mutants displayed on yeast. Data are the mean ± SD of n = 3 individual yeast clones, normalized to MFI from binding to 47_WT_. (e) Binding of B6H12, CV-1, hSIRPα, and mSIRPα to yeast displayed 47_WT_, 47_A30P_, and 47_Q31P_ variants. Data are the mean ± SD of n = 3 individual yeast clones, normalized to MFI from binding to 47_WT_. (f) Crystal structures of CD47 (red) binding SIRPα (dark pink, left) [PDB: 2JJS] and B6H12 (light blue, right) [PDB: 5TZU], identifying residues A30 (gold) and Q31 (blue). Lower panels are enlargements of the boxed regions in the full structures. (g) Binding of B6H12, TJC4, Hu5F9, and hSIRPα to yeast displayed 47_WT_, 47_A30P_, and 47_Q31P_ variants. Data are the mean ± SD of n = 3 individual yeast clones, normalized to MFI from binding to 47_WT_. (h) Binding of B6H12, TJC4, hSIRPα, and mSIRPα to full-length 47_WT_, 47_A30P_, and 47_Q31P_ expressed on primary human T cells. Data are the mean ± SD of n = 3 donors, normalized to fraction binding to 47_WT_. (i) Binding of B6H12, hSIRPα, and mSIRPα to full-length 47_WT_, 47_A30P_, and 47_Q31P_ expressed on Jurkat cells with endogenous 47_KO_. Data are the mean ± SD of n = 3 independent experiments, normalized to fraction binding to 47_WT_. (j) Quantification of phagocytosis by primary human macrophages of CSFE labeled Jurkats with endogenous 47_KO_, expressing 47_WT_, 47_A30P_, or 47_Q31P_ variants. Data are the mean ± SD of triplicate wells (n = 3). Data is reproducible across four different macrophage donors. (k – l) T cell depletion model. (k) T cell depletion scheme. Non-tumor bearing mice were treated IV with 5⨉10^6^ 47_WT_- or 47_E_-CD19.28ζ-nLuc-CAR T cells (with endogenous 47_KO_), ± two doses of B6H12 (250 µg/dose; IP) on days 3 and 5 post CAR T treatment. Mice were imaged by BLI before (day 3) and after (day 7) αCD47 treatment, and had blood drawn on day 6 for detection of T cells. (l) Quantification of CD8^+^ T cells treated as described in (k) in the blood after B6H12 treatment (day 6). Data are the mean ± SD of n = 5 mice. 47_WT_ is a shared condition duplicated in T cell depletion in Extended Data Fig. 2i. [(d), (e), (f), (g), (i), (j), and (l)] Two-way ANOVA test with Tukey’s multiple comparison test. ns = not significant. (g), (h) and (i) comparison is between indicated group and binding to 47_WT_ expressing cells. See also Extended Data Figures 4 and 5.

To assess binding to wild-type (WT) SIRPα, we next displayed 47_A30P_ and 47_Q31P_ mutants on yeast in a more natural orientation, with a free CD47 N-terminus^57^ (**Extended Data Fig. 4h,i**). Both mutants retained binding to mouse and human SIRPα but demonstrated no binding to B6H12, even at the high concentration of 3 µM (**Fig. 5e**). These results are consistent with current structural understanding of CD47-SIRPα interactions, whereby SIRPα predominantly contacts CD47 through residues in the CD47 FG loop and N-terminus^58^ (**Fig. 5f**), while forming more minor contacts with the CD47 BC loop, which encompasses T26 – Q31^58^ (**Fig. 5f, Extended Data Fig. 4j**), potentially leaving residues in this region amenable to mutation. Because the CD47 BC loop lies near the critical CD47 FG loop, it can serve as an anchoring point for αCD47 blocking mAbs like B6H12^59, 60^, with the Q31 residue appearing particularly important for antibody binding^59, 60^ (**Fig. 5f**).

To determine the potential for the CD47 mutants to evade binding by CD47 blocking mAbs currently in clinical trials, we analyzed binding of TJC4^61, 62^ (lemzoparlimab; Phase II) and Hu5F9^63^ (magrolimab; Phase III) to yeast displayed CD47 mutants. We began by performing an alanine scan of the entire BC loop (T26 – Q31), comparing binding to human SIRPα, B6H12, TJC4, and Hu5F9 (**Fig. 5g**, **Extended Data Fig. 4k**). Most mutations allowed for some retained SIRPα binding, with mutations to A30 or Q31 manifesting the most minimal impact on SIRPα binding. Hu5F9, which has a binding footprint that largely overlaps with SIRPα^64^, demonstrated minimal loss of binding to any of the BC loop mutants, including 47_A30P_ or 47_Q31P_. However, TJC4, which structurally binds CD47 similarly to B6H12^65^, no longer bound 47_A30P_, 47_Q31P_, and 47_A30P-Q31A_, nor did it bind 47_E29A_, which manifests an additional mutation that did not affect B6H12 binding (**Fig. 5g**, **Extended Data Fig. 4k**). We next profiled binding of SIRPα, B6H12 and TJC4 to full-length 47_WT_, 47_A30P_, 47_Q31P_, and 47_A30P-Q31A_, expressed on primary human T cells. Binding of both human and mouse SIRPα was largely unaffected by any of the three mutants indicating that these variants are predicted to retain their function as “don’t eat me” signals (**Fig. 5h**, **Extended Data Fig. 5a**). However, we detected no B6H12 binding over baseline to any of the three mutants, and TJC4 binding was completely ablated by the 47_A30P-Q31A_ double mutant. These data demonstrate that CD47 mutations to the BC loop, and A30 and Q31 specifically, generate proteins that retain SIRPα binding but are exempt from binding to multiple CD47 mAbs, providing proof-of-concept for the ability to engineer CD47 variants predicted to protect T cells from phagocytosis induced by αCD47 mAbs, and thereby induce tumor specific phagocytosis while sparing T cells in the TME.

### Expression of engineered CD47 protects T cells from αCD47 induced phagocytosis

We next measured phagocytosis of 47_KO_ Jurkat cells engineered to express either 47_WT_, 47_A30P_, or 47_Q31P_ by human donor macrophages. CD47 mutants expressed on Jurkats demonstrated similar binding properties to αCD47 mAbs and SIRPα as observed on primary T cells, with 47_Q31P_ leading to the greatest loss of B6H12 binding (**Fig. 5i**, **Extended Data Fig. 5b**). Across multiple donors, we observed that expression of either mutant significantly reduced phagocytosis after B6H12 incubation (**Fig. 5j, Extended Data Fig. 5c,d**), but 47_A30P_ provided less protection compared to 47_Q31P_, which completely prevented additional phagocytosis after incubation with B6H12. Based on this promising profile, we chose to move forward with 47_Q31P_, referred to as 47_E_ (“engineered CD47”), for further study.

We interrogated whether 47_E_ expression by T cells could protect against B6H12 mediated depletion *in vivo*. To prevent B6H12 binding to endogenous CD47, we knocked out endogenous CD47 using CRISPR/Cas9, before introducing CAR, TCR, and/or 47_E_ or wild-type CD47 (47_WT_). Following administration of 47_WT_- or 47_E_-CD19.28ζ-nLuc CAR T cells to non-tumor bearing mice we observed similar levels of T cells as measured by BLI across groups prior to B6H12 administration (**Fig. 5k**, **Extended Data Fig. 5e,f**), but 47_WT_-CAR T were completely depleted *in vivo* after B6H12 treatment as previously shown (**Fig. 5l**, **Extended Data Fig. 5e-g**). In contrast, 47_E_-CAR T were not depleted after B6H12 administration and persisted at similar levels compared to mice that had only received PBS vehicle control (**Figure 5l, Extended Data Figure 5e-g**). These data demonstrate that 47_E_ functions as a “don’t eat me” signal *in vivo*, but remains inert from binding B6H12, and thereby protects T cells from phagocytosis after CD47 blockade.

### CAR T cell treatment leads to macrophage tumor infiltration which is augmented in animals treated with 47_E_-CAR T cells plus αCD47

To investigate the effects of combining 47_E_-CAR T with αCD47 on the TME, we treated mice bearing orthotopically implanted 143B osteosarcoma tumors with no T cells, mock, 47_WT_- or 47_E_-Her2.BBζ-CAR T ± B6H12 (**Fig. 6a**, **Extended Data Fig. 6a,b**). After 8 days, tumors from CAR T recipients demonstrated significant increases in F4/80^+^ murine macrophages, compared with mock or untreated animals (**Fig. 6a-c**). B6H12 treatment did not significantly impact macrophage levels in the TME of 47_E_-CAR T recipients, whereas macrophages were substantially reduced following B6H12 therapy in 47_WT_-CAR T recipients (**Fig. 6b-d**, **Extended Data Fig. 6c-g**). CAR T and macrophage infiltration into tumors were highly correlated, consistent with a model whereby CAR T cells recruit macrophages into the tumor, and macrophage persistence is dependent upon CAR T persistence in the tumor (**Fig. 6d**, **Extended Data Fig. 6e-g**). ScRNA-seq analysis profiling of both human (tumor and T cells) and mouse (immune and fibroblast) cells in dissociated tumors (**Extended Data Fig. 7a,b**) confirmed CAR T mediated increases in the frequencies of macrophages within the tumor, which was ablated in 47_WT_-CAR T recipients following B6H12 treatment, but persisted in 47_E_-CAR T recipients (**Fig. 6e,f**).

**Figure 6:**
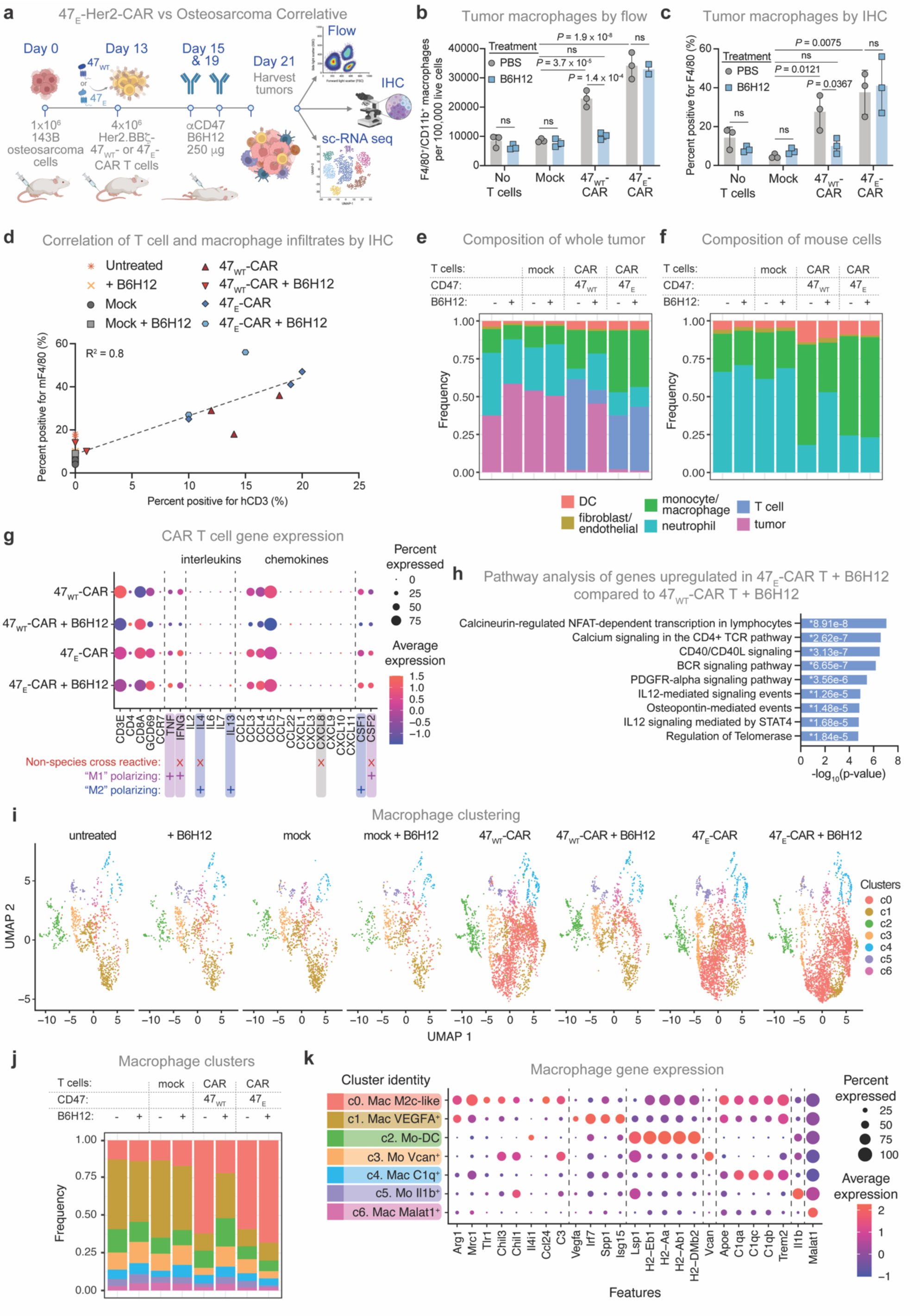
Paired CAR T and αCD47 treatment leads to recruitment of distinct macrophage populations to the tumor microenvironment. (a) Scheme of mechanistic study in 143B osteosarcoma. Mice were engrafted orthotopically in the tibia periosteum with 1⨉10^6^ 143B cells and treated IV with no T cells, or 4⨉10^6^ mock, 47_WT_- or 47_E_-Her2.BBζ-CAR T cells (with endogenous 47_KO_) on day 13. Mice were then treated ± two doses of B6H12 (250 µg/dose; IP) on days 15 and 19. Tumors were excised on day 21 and then analyzed via flow cytometry, IHC, and scRNA-seq. (b) Quantification of mF4/80^+^/mCD11b^+^ macrophages identified by flow cytometry of dissociated tumors treated as described in (a). Data are the mean ± SD of n = 2 (47_E_-CAR + B6H12) or n = 3 (all other samples) mice. (c) Quantification of mF4/80 staining of IHC sections produced from tumor sections generated as described in (a). Data are the mean ± SD of n = 3 mice. (d) Correlation of quantification of hCD3 and mF4/80 staining in IHC sections of tumors treated as described in (a). Data points are representative of individual tumors, colored by treatment group (n = 23). R^2^ calculated by simple linear regression. (e and f) Composition of (e) all cell types or (f) solely mouse cells identified via scRNA-seq from tumors treated as described in (a). Data are derived from n = 53,062 cells pooled from 8 experimental conditions with three mice per treatment group. Cell types assigned using SingleR automated cell type recognition. (g) Dot Plot depicting scRNA-seq expression of selected T cell subset markers, cytokines, and chemokines. n = 11,044 human tumor infiltrating T cells from 4 experimental conditions described in (a). Genes encoding proteins involved in macrophage “M1” or “M2” polarization are indicated by purple or blue plus signs, respectively. Genes encoding proteins that are non-species cross reactive between human and mouse are marked with a red “x.” (h) Enrichr pathway analysis of the top 100 upregulated genes in tumor infiltrating CAR T cells in 47_E_-CAR + B6H12 treated tumors compared with 47_WT_-CAR + B6H12 treated tumors. The NCI-Nature 2016 collection was queried with gene IDs. (i) UMAPs of the identified macrophage/monocyte population [Extended Data Fig. 7a], subsetted and re-clustered, colored by cluster. UMAPs represent distinct treatment conditions. Dots represent individual cells. *n* = 13,082 cells from 8 experimental conditions described in (a). (j) Composition of macrophage clusters identified in (i) across experimental conditions described in (a). (k) Dot plot depicting scRNA-seq expression of selected cluster-defining genes within the macrophage populations identified in (i). P adj. < 0.0001 for each selected gene. [(b) and (c)] Two-way ANOVA test with Tukey’s multiple comparison test. ns = not significant. See also Extended Data Figure 6 and 7 and Supplementary Table 3.

We next profiled the TME to identify genes potentially responsible for macrophage recruitment and activation. 47_WT_- and 47_E_-CAR T recipients demonstrated robust T cell expression of TNFα, IFN γ, CCL3, CCL4, and CCL5, CSF1 (M-SCF) and CSF2 (GM-CSF) (**Fig. 6g**), which collectively attract and activate monocytes and macrophages^66, 67^ and have been shown to be responsible for T cell mediated macrophage recruitment into tumors^68^. Of note, because IFNγ and CSF2, are not species cross-reactive^69^, the full potential of T cell – macrophage crosstalk in this model is likely underestimated. 47_E_-CAR T gene expression was largely unaffected by treatment with B6H12, however T cells in the TME of B6H12 recipients co-treated with 47_E_-CAR T showed 595 differentially expressed genes (DEGs) compared to those co-treated with 47_WT_-CAR T (**Extended Data Fig. 7c**, **Supplementary Table 1**), including gene sets associated with IL-12 signaling^70^ and CD40/CD40L signaling^71, 72^ (**Fig. 6h**), consistent with increased crosstalk between myeloid cells and T cells within the 47_E_-CAR T TME compared to that found in recipients of 47_WT_-CAR T after B6H12 co-treatment. Together, these data reveal potent effects of CAR T on macrophage recruitment into the TME and demonstrate sustained increases in both CAR T cells and macrophages within the TME in 47_E_-CAR T plus B6H12 recipients, which is associated with enhanced CAR T signaling of numerous pathways associated with improved functionality.

DEG analyses in the major macrophage cluster across treatments showed that treatment with 47_WT_-CAR T alone induced 621 DEGs in macrophages compared to the untreated condition, and this effect was magnified following 47_E_-CAR T co-treatment with B6H12, with 718 DEGs (**Extended Data Fig. 7d**, **Supplementary Table 2**). Interestingly, the effect of B6H12 therapy on macrophage gene expression when administered as a single agent was minimal (46 DEGs) (**Extended Data Fig. 7d**). However, B6H12 co-administration with 47_WT_-CAR T dramatically reduced macrophage DEGs, likely due to CAR T depletion (**Extended Data Fig. 7d**). Pathway analysis of genes upregulated by 47_E_-CAR T plus B6H12 highlighted macrophage activation indicated by enrichment of lysosome, complement, antigen presentation, and phagosome pathways^73, 74^ (**Extended Data Fig. 7e**). Together, these results demonstrate a feed forward loop wherein CAR T cells drive recruitment and activation of macrophages within the TME and simultaneously, macrophages enhance activating pathways within the 47_E_-CAR T TME, but not within the 47_WT_-CAR T TME, wherein CAR T depletion abrogates the cycle after CD47 blockade.

To further characterize changes in the macrophage compartment induced by 47_E_-CAR T, we re-clustered the macrophage/monocyte cluster (**Fig. 6i**, **Extended Data Fig. 7f**). As observed in clinical data (**Fig. 3f,g**), we identified numerous macrophages that contained hCD3ε mRNA within multiple macrophage sub-clusters (**Extended Data Fig. 7g**), consistent with macrophage mediated phagocytosis of T cells. We also observed expansion of macrophage cluster “c0,” which was enriched following CAR T treatment and further expanded following 47_E_-CAR T plus B6H12, but nearly completely absent after 47_WT_-CAR T plus B6H12 (**Fig. 6i,j**), suggesting that these macrophages are dependent upon CAR T cell accumulation within the TME. Key DEGs in the expanded cluster were associated with M2c-like macrophages^75^ (**Fig. 6j,k**, **Supplementary Table 3**). The M2c-like cluster (c0) highly expressed canonical M2 macrophage genes such as *Arg1*, *Mrc1*, and *Chil3*, in addition to M2c genes such as *Tlr1*^75^ (**Fig. 6k**). While M2 macrophages are generally understood to be pro-tumorigenic, they have also been demonstrated to manifest strong phagocytic potential, especially those in M2c subclass^76, 77^. Together, these data demonstrate robust crosstalk between CAR T cells and macrophages in tumors, with significant dependency of the macrophage population on CAR T persistence in the TME, and macrophage induced induction of gene expression programs in the CAR T predicted to enhance anti-tumor effects.

### 47_E_-CAR T plus αCD47 induces enhanced antitumor efficacy

We next assessed the antitumor effects of combined 47_WT_-vs 47_E_-T cells plus αCD47 therapy in multiple tumor models treated with cell therapies. We first administered NY-ESO-1-TCR-Antares T cells to mice with A375 melanoma flank xenografts (**Fig. 7a**). While 47 _WT_-NY-ESO-1 T cells were depleted after B6H12 treatment, 47_E_-NY-ESO-1 T cells were protected (**Fig. 1e**, **Fig. 7b,c**, **Extended Data Fig. 8a,b**). A375 tumor growth was minimally slowed by B6H12 plus mock T cells (**Fig. 7d**) and treatment with a low dose of 1⨉10^6^ 47_E_-NY-ESO-1 T cells alone led to initial tumor control, but ultimate tumor outgrowth in 4/5 mice treated (**Fig. 7d**). By contrast, mice treated with 47_E_-NY-ESO- 1 T cells and B6H12 demonstrated complete tumor control and cure in 5/5 mice treated (**Fig. 7d**), and these results were confirmed using a second donor (**Extended Data Fig. 8c**). We next interrogated pairing B6H12 with 47_E_-CAR T in a metastatic neuroblastoma model. Metastatic CHLA-255-fLuc bearing mice received 47_WT_- or 47_E_-B7H3.BBζ-nLuc-CAR T cells, ± B6H12, and we observed persistence of 47_E_-CAR T, but not 47_WT_-CAR T (**Extended Data Fig. 8d-g**). When B6H12 was administered, 47_WT_-CAR T mediated no improvement in anti-tumor efficacy, while 47_E_-CAR T mediated significantly improved antitumor efficacy, compared to either agent alone (**Extended Data Fig. 8g,h**).

**Figure 7:**
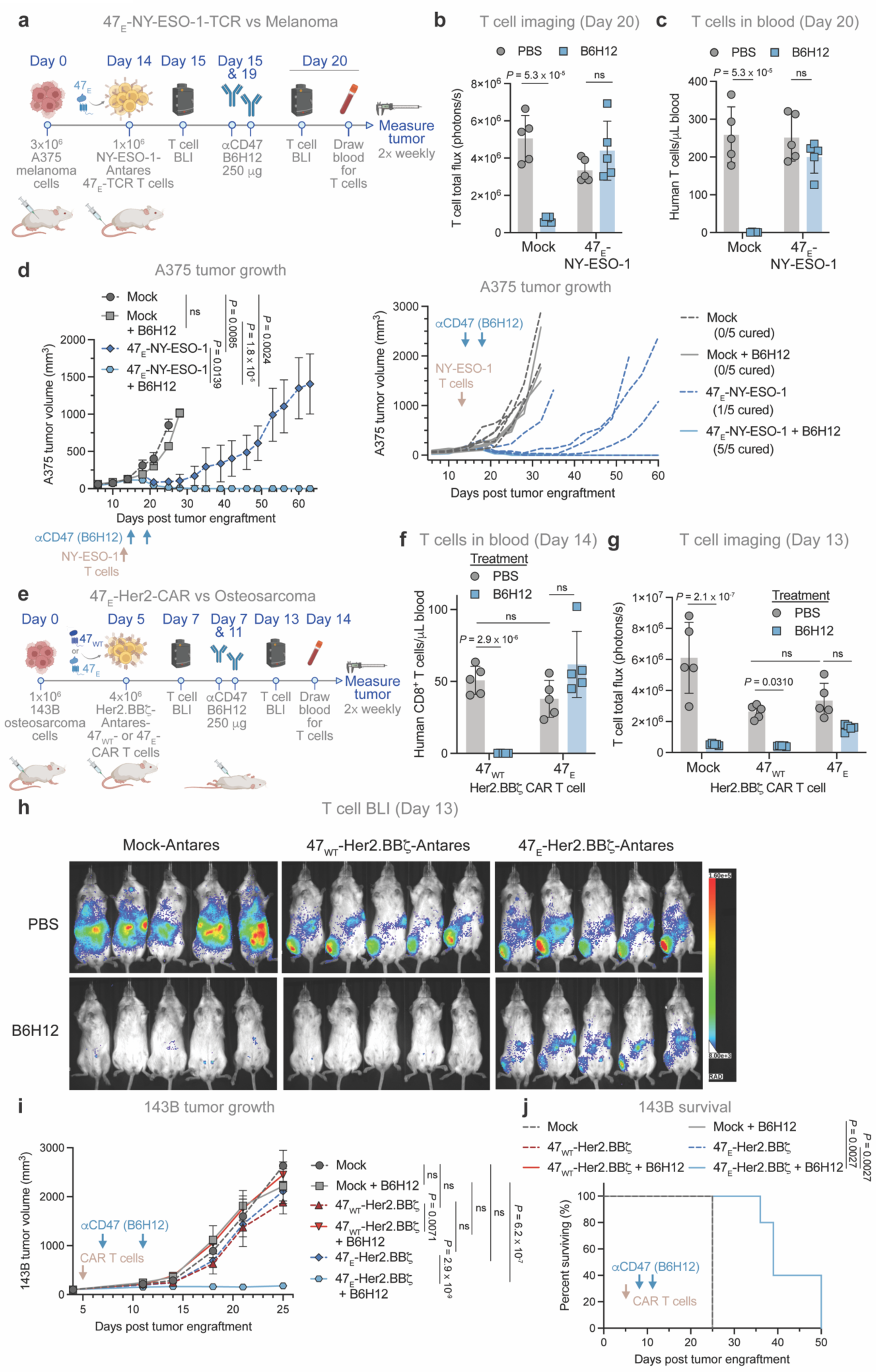
Expression of engineered CD47 on T cells permits pairing with anti-CD47 therapy. (a – d) A375 melanoma model (a) A375 treatment scheme with NY-ESO-1-TCR T cells. Mice were engrafted SQ with 3⨉10^6^ A375 cells and treated IV with 1⨉10^6^ mock-Antares or 47_E_-NY-ESO-1-Antares-TCR T cells (with endogenous 47_KO_) on day 14. Mice were then treated ± two doses of B6H12 (250 µg/dose; IP) on days 15 and 19. (b and c) Quantification of T cells by (b) BLI and (c) in the blood (hCD45^+^) in the A375 – NY-ESO-1 model. Mice engrafted SQ with 3⨉10^6^ A375 were treated IV with 1⨉10^6^ mock-Antares or 47_E_-NY-ESO-1-Antares-TCR T cells on day 7 ± three doses of B6H12 (250 µg/dose; IP) on days 9, 11, and 14. Mice were imaged by BLI before (day 9) and after (day 14) αCD47 treatment. Blood was collected on day 15. Data are the mean ± SD of n = 5 mice. Mock is a shared control duplicated in T cell depletion in Fig. 1e. (d) A375 tumor growth treated as described in (a). Data are the mean ± SEM (left) or individual tumor growth traces (right) of n = 5 mice/arm. (e – f) 143B osteosarcoma model (e) 143B treatment scheme. Mice were engrafted orthotopically in the tibia periosteum with 1⨉10^6^ 143B cells and treated IV with 4⨉10^6^ mock-Antares, or 47_WT_- or 47_E_-Her2.BBζ-Antares-CAR T cells (with endogenous 47_KO_) on day 5. Mice were then treated ± two doses of B6H12 (250 µg/dose; IP) on days 7 and 11. Mice were imaged by BLI before (day 7) and after (day 13) αCD47 treatment, and had blood drawn on day 14. (f and g) Quantification of T cells (f) in the blood (hCD8^+^, day 14) and by (g) BLI (day 13) in the 143B tumor model treated as described in (e). Data are the mean ± SD of n = 5 mice. (h) T cell BLI in the 143B model on day 13, treated as described in (e). Mice are imaged on their backs; 143B tumors are engrafted in the right leg of each mouse. (i and j) 143B tumor (i) growth and (j) survival, treated as described in (e). Data are (i) the mean ± SEM or (j) representative of n = 5 mice/arm. [(b) and (c)] Unpaired two-tailed Student’s *t* test. ns = not significant. [(d), (f), (g), and (i)] Two-way ANOVA test with Tukey’s multiple comparison test. ns = not significant. [(j)] Log-rank Mantel-Cox test. See also Extended Data Figure 8.

Finally, we explored the antitumor effects of combination therapy in the aggressive orthotopic osteosarcoma model, 143B, where both CAR T and αCD47 therapy have minimal effect as monotherapies, but where we had observed striking T cell mediated recruitment of macrophages into tumors. Mice bearing 143B tumors in the tibia periosteum received 47_WT_- or 47_E_-Her2.BBζ-Antares-CAR T cells, ± B6H12 (**Fig. 7e**). Following B6H12 administration, 47_WT_-CAR T were completely depleted, while 47_E_-CAR T persisted over the course of multiple weeks (**Fig. 7f-h**, **Extended Data Fig. 8i-l**). Over two independent experiments, we observed no meaningful tumor control after treatment with 47_WT_-CAR T or 47_E_-CAR T alone, or mock T cells paired with B6H12 (**Fig. 7i,j**, **Extended Data Fig. 8m,n**) and B6H12 combined with 47_WT_-CAR T delivered no significant anti-tumor efficacy (**Fig. 7i,j**, **Extended Data Fig. 8m,n**). However, strikingly, B6H12 plus 47_E_-CAR T induced marked tumor control, a significant delay in tumor outgrowth, and improvement in overall survival (**Fig. 7i,j**, **Extended Data Fig. 8m,n**). These results demonstrate strong synergy in solid and metastatic tumors using CD47 blockade paired with 47_E_ expressing therapeutic T cells, even at low doses and in conditions where both single-agent therapies showed no activity. Together these data illustrate that protection of CAR T cells from macrophage mediated phagocytosis results in a dramatic and sustained influx of macrophages within the TME, associated T cell-macrophage crosstalk, and remarkably enhanced antitumor efficacy compared to treatment with either agent alone.

## Discussion

The data presented here demonstrate a critical role for the CD47-SIRPα axis in modulating T cell survival *in vivo*. Previous studies reported that CD47 is necessary to prevent immune rejection^78^ and that CD47 overexpression, when combined with MHC knockout, imparts resistance of CAR T cells, pancreatic islets, and iPSCs to allogeneic immune rejection^79–81^. Our findings demonstrate that even in the absence of immune rejection, CD47 is required for survival of adoptively transferred T cells and CD47 overexpression improves CAR T cell persistence and efficacy (**Fig. 2e,f**). These results align with recently published data demonstrating that CD47_KO_ CAR T cells show limited persistence in xenograft models^82^ and previous murine studies demonstrating reduced numbers of total and antigen specific lymphocytes, increased susceptibility to infection, and reduced susceptibility to autoimmunity in CD47_KO_ and SIRPα_KO_ mice^83–85^. Relevance to the clinical setting is provided by the observation of lymphopenia in studies of magrolimab, an αCD47 mAb, and evorpacept, a CV-1-Fc fusion protein, which blocks CD47 engagement of SIRPα^17, 18^. We also observed that αCD47 treatment of NSG murine hosts induced rapid and complete macrophage mediated depletion of adoptively transferred T cells, ablating antitumor efficacy (**Fig. 1**). Remarkably, the depletion is sufficiently rapid and potent to mediate complete protection in a lethal CAR T cell toxicity model (**Fig. 4**), thus providing data to support testing of CD47/SIRPα blockers to mitigate toxicities resulting from adoptive T cell therapeutics, which could have immediate clinical benefits.

Our data further provide evidence that macrophage phagocytosis limits persistence of adoptively transferred T cells even in the absence of CD47/SIRPα blockade, as scRNA-seq revealed CAR T cell RNA in myeloid cells of mice that were not treated with CD47 blockade (**Extended Data Fig. 7g**), and we observed improved T cell persistence following macrophage depletion (**Fig. 2i,j**). Clinical scRNA-seq data also consistently identified myeloid cells possessing CAR genes (**Fig. 3f,g**). Together, the findings are consistent with a model wherein macrophage phagocytosis plays a significant, and previously unrecognized role, in regulating T cell homeostasis. Future work is needed to define the relationship between T cell states, expression of pro-phagocytic and anti-phagocytic receptors, and phagocytosis susceptibility. It is well recognized that a major toxicity of αCD47 therapy is anemia, occurring as a result of depletion of aged red blood cells^15–17, 86^. We observed increased expression of “pro-eat me” signals and decreased expression of CD47 on T cells following prolonged culture (**Fig. 3c,d**), suggesting that the CD47-SIRPα axis may similarly regulate clearance of aged T cells.

Tumor associated macrophages (TAMs) are among the most plentiful cells in the TME, and there has been great interest in harnessing their antitumor properties, but the optimal approach has remained elusive^1–3, 87^. CSF1R and CCR2 inhibitors inhibit recruitment of monocytes to the TME and reduce or eliminate TAMs^7–9^ and have been tested in dozens of clinical trials, but thus far significant antitumor effects has not been demonstrated^2, 3^. These results have led investigators to pursue more targeted approaches to modulate macrophage states from those with immunosuppressive profiles, including the M2-like subset, toward those with more inflammatory profile, but thus far these approaches have also had limited clinical success^2, 3, 88^.

Systemic blockade of the CD47/SIRPα axis augments macrophage phagocytosis broadly and mediates antitumor effects in several preclinical models^12–14, 36, 47, 89, 90^. In the clinic, these agents have demonstrated efficacy when combined with tumor specific mAbs in non-Hodgkin lymphoma^15^ and with epigenetic modifiers in AML^16^, however clinical benefit as single agents has been limited, and significant activity in solid cancers has not been reported^18, 91^. Several creative strategies are under development to augment the safety and potency of αCD47 therapy, most of which have focused on more effective delivery to the TME^92–96^, including delivery of αCD47 peptides with CAR T cells^97–100^, however the therapeutic benefit from these preclinical studies has been modest thus far. The data presented here suggest that this conundrum may be explained by the double-edged sword that TAMs represent within the TME, whereby antitumor effects resulting from maneuvers that augment macrophage phagocytosis are offset by phagocytosis of tumor reactive T cells. Conversely, eliminating or reducing TAMs may diminish immunosuppression and phagocytosis of tumor infiltrating T cells, but these benefits are predicted to be offset by loss of tumor phagocytosis by macrophages. Here we demonstrate dramatically augmented antitumor effects using an approach which simultaneously enhances macrophage phagocytosis of tumor cells while protecting tumor reactive T cells from phagocytosis. Our data demonstrate that pairing 47_E_-CAR T therapy with αCD47 is an exciting prospect that could enhance the potency of adoptive T cell therapies, especially in solid cancers (**Fig. 7**).

We observed that CAR T cells induce a rapid influx of macrophages into tumors (**Fig. 6**), which is dependent upon CAR T cell persistence, since treatment with αCD47, which depletes 47_WT_-CAR T also dampened TAM infiltration. In contrast, high TAM levels were sustained in animals receiving αCD47 with 47_E_-CAR T. In 47_E_-CAR T recipients, we further saw evidence for extensive crosstalk between myeloid cells and T cells in the TME, as evidenced by the induction of IL-12, CD40L and NFκB signaling in T cells (**Fig. 6h**), all of which have been demonstrated to augment antitumor effects^101–107^. 47_E_ protected T cells from αCD47 mediated phagocytosis (**Fig. 5j-l**), and allowed for the beneficial effects of CD47 therapy, such as tumor phagocytosis, antigen presentation, and a pro-inflammatory TME^108^, without sacrificing T cells engaged in the same antitumor effort. Additional studies are needed to profile the pairing of αCD47 and CAR T therapy in fully immunocompetent systems, as it is critical to understand the impact of other immune cell types, including endogenous T and NK cells^109^ on this phenomenon. We demonstrated striking synergy when pairing αCD47 antibody B6H12 with adoptive T cell therapy expressing 47_E_, mediated through recruitment of tumor macrophages (**Fig. 6 & 7**). While B6H12 is a research grade antibody^36–39^, we found that the CD47 variants generated in this study also ablated binding to the clinical grade antibody TJC4 (lemzoparlimab)^61, 62, 65^ (**Fig. 5g-i**). Thus, these mutations could allow pairing of CAR T therapy with clinical grade αCD47 antibodies. Further enhancements could be developed for this approach, as innumerable engineering strategies have been implemented to improve the efficacy of CAR T cells ^31, 35, 110–114^, while CD47 blockade is often most efficacious when paired with additional antibodies ^13, 15, 100^. Further, while we chose to knock out endogenous CD47 and virally transduce in 47_E_, potentially deriving benefits from 47_OE_ (**Fig. 2e,f**)^45^, it may be possible to introduce 47_E_ at the endogenous CD47 locus in the future using a base-editor^115, 116^ or prime-editor^117^. Multiple studies have reported on CAR T cells secreting αCD47 agents^97–100^. These data suggest that future iterations of this approach could make use of 47_E_ expressing CAR T cells secreting αCD47 proteins, packaging both therapies into a single “living” drug.

Immunotherapy has led to a revolution in cancer patient treatment and survival. However, most immunotherapies, including adoptive T cell therapy and CD47 blockade have demonstrated limited efficacy in the treatment of solid tumors to date. By allowing for the combination of these two different immunotherapies, our work provides a potential pathway to a novel kind of immunotherapy that takes advantage of the power of both the innate and adaptive immune systems to deliver exquisite tumor control.

## Acknowledgements

This work was supported by National Institutes of Health grants 1 R01 CA263500-01 (C.L.M. and M.M.); a EPICC Translational Research Grant (St. Baldrick’s Foundation, C.L.M.); Parker Institute for Cancer Immunotherapy (C.L.M.); the Virginia and D.K. Ludwig Fund for Cancer Research (C.L.M.), and an NCI grant R35 CA220434 (I.L.W). C.L.M., R.G.M., L.L., and Z.G. are members of the Parker Institute for Cancer Immunotherapy, which supports the Stanford University Cancer Immunotherapy Program. A.L. is supported by the Nuovo-Soldati Foundation and by ITMO Cancer AVIESAN (Alliance Nationale pour les Sciences de la Vie et de la Santé/National Alliance for the Life Sciences and Health) within the framework of the French Cancer Plan. R.G.M. is the Taube Distinguished Scholar for Pediatric Immunotherapy at Stanford University School of Medicine. B.J.M. was supported by a Stanford Interdisciplinary Graduate Fellowship. K.A.F. was supported by the National Science Foundation Graduate Research Fellowship under grant DGE-1656518. S.H. was supported by a U54 CA232568-01 grant. O.K. was supported by a Paul and Daisy Soros Fellowship for New Americans. Sorting was performed on an instrument in the Stanford Shared FACS Facility obtained using NIH S10 Shared Instrument Grant S10RR025518-01. The authors acknowledge the Stanford Shared FACS facility for sorting tumor samples, the Stanford Human Pathology/Histology Service Center for immunohistochemistry staining, the Stanford Veterinary Service Center for general support of mouse work and necropsy studies, and ALX Oncology for contributing the CV-1 reagent. Illustrations were created with BioRender.com.

## Author Contributions

S.A.Y.-H., J.T., and C.L.M. conceived the idea of the study and designed experiments. S.A.Y.-H., J.T., B.J.M., K.A.F., M.T.R., A.L., S.D., N.M.-V., P.X., A.D., M.H.D., L.L., C.W.M., Y.C., S.H., K.D.M., A.B., O.K., J.H., S.L.W., S.F.-P., and R.G.M. performed experiments and acquired data. S.A.Y.-H., J.T., B.J.M., K.A.F, M.T.R, A.L., S.D., N.M.-V., A.D., Z.G., J.Y.S., S.F.-P., R.G.M., E.S., J.R.C., and C.L.M. analyzed data and interpreted results. P.H.S., M.M., I.L.W., B.S., E.S., J.R.C., and C.L.M. supervised the work. S.A.Y.-H., E.S., and C.L.M. wrote the manuscript with feedback from all authors.

## Materials and Methods

### Cell lines

The Nalm6 B-ALL cell line was provided by David Barrett (Children’s Hospital of Philadelphia) and retrovirally transduced to express GFP and firefly luciferase. 143B osteosarcoma cells (ATCC) were retrovirally transduced with human CD19. CHLA-255 neuroblastoma line was provided by Robert Seeger (Children’s Hospital Los Angeles) and retrovirally transduced with GFP and firefly luciferase. MG63.3 was provided by Chand Khanna (National Cancer Institute, National Institutes of Health) and retrovirally transduced with GFP and firefly luciferase. D425 was provided by S. Chesier (Stanford University, Stanford, CA) and retrovirally transduced to express GFP and firefly luciferase. A375 melanoma cells were obtained from ATCC. The 293GP retroviral packaging line was provided by the Surgery Branch (National Cancer Institute, National Institutes of Health). Expi293 protein production cells were obtained from ATCC. D425 cells were maintained in serum-free media supplemented with B27 (Thermo Fisher Scientific), EGF, FGF (Shenandoah Biotechnology), human recombinant LIF (Millipore), and Heparin (StemCell Technologies). Nalm6, 143B, A375, MG63.3, and CHLA-255 were cultured in RPMI-1640 (Gibco). 293GP were cultured in DMEM (Gibco). Expi293 cells were cultured in Expi293 media (Thermo Fisher Scientific). Cell line culture media was supplemented with 10% FBS, 10mM HEPES, 2mM L-glutamine, 100 U/mL penicillin, and 100μg/mL streptomycin (Gibco), except for Expi293 media. STR DNA profiling of all cell lines was conducted once per year (Genetica Cell Line testing). All cell lines were routinely tested for mycoplasma. Cell lines were cultured at 37°C in a 5% CO_2_ environment.

### Source of primary human T cells and macrophages

Buffy coats from healthy donors were purchased from the Stanford Blood Center under an IRB-exempt-protocol. Primary human T cells were purified by negative selection using the RosetteSep Human T cell Enrichment kit (Stem Cell Technologies) and SepMate-50 tubes. T cells were cryopreserved at 2×10^7^ cells per mL in CryoStor CS10 cryopreservation media (Stem Cell Technologies) until use. Primary peripheral monocytes were purified through successive density gradients using Ficoll (Sigma-Aldrich) and Percoll (GE Healthcare). Monocytes were then differentiated into macrophages by 7–9 d of culture in IMDM + 10% AB human serum (Life Technologies).

### Viral vector construction

All retroviral constructs were cloned into the MSGV1 retroviral vector^118^. B7H3.BBζ was generated by fusing, from N to C terminus, a human GM-CSF leader sequence, scFv derived from MGA271 in the VH-VL orientation and (GGGS)_3_ linker sequence, CD8α hinge and transmembrane sequence, and human 4-1BB and CD3ζ intracellular signaling domains. GD2.BBζ, Her2.BBζ, and CD19.BBζ were generated by cloning scFvs derived from 14G2A, 4D5, and FMC63 antibodies, respectively into the B7H3.BBζ vector. CD19.28ζ was generated by replacing the 4-1BB domain in CD19.BBζ with the intracellular signaling domain of human CD28. PIP.28ζ and PIP.BBζ were generated by replacing the FMC63 scFv with the 2.5F knottin^50^ in the CD19.28ζ and CD19.BBζ vectors, respectively. The *in vivo* T cell activation reporter was constructed by cloning a sequence containing firefly luciferase into the pGreenFire1-NF-κB lentiviral vector (System Biosciences) under the NF-κB responsive promoter. CD47 vectors were generated by inserting codon-optimized CD47 sequences (mutant and wild-type) in place of the CD19.BBζ sequence. For *in vivo* tracking, CAR-nLuc plasmids were generated by replacing the stop codon in the CD3ζ with a sequence containing a porcine teschovirus-1 2A (P2A) ribosomal skipping sequence, followed by nanoluciferase. The NY-ESO-1 TCR construct was generated by inserting the NY-ESO-1 α chain, followed by a P2A sequence, followed by the β chain in place of CD19.BBζ.

### Virus production

Retroviral supernatant was packaged using 293GP cells and the RD114 envelope plasmid. In brief, 11μg RD114 and 22μg of the corresponding MSGV1 transfer plasmid were delivered to 293GP cells grown on 150mm poly-D-lysine dishes (Corning) to 80% confluency by transient transfection with Lipofectamine 2000 (Thermo Fisher). Media was replenished every 24 hours. Virus production was performed side-by-side for comparable CAR, TCR, and CD47 constructs. Retroviral supernatant was harvested 48 and 72-hour post transfection. Supernatant from replicate dishes were pooled, centrifuged to deplete cell debris, and stored at −80C until use. Third-generation, self-inactivating lentiviral supernatant was similarly produced with 293T cells using 7μg pMD2.G (VSVg) envelope, 18μg pMDLg/pRRE (Gag/Pol), 18μg pRSV-Rev, and 20μg the corresponding transfer plasmids.

### CAR T and TCR T manufacturing

At Day 0, primary human T cells were thawed and activated with anti-CD3/CD28 Human T-Expander Dynabeads (Thermo Fisher) at a 3:1 bead to cell ratio. On Day 2 virus coated culture plates were prepared on non-TC-treated 12-well plates that had been pre-coated with RetroNectin (Takara Bio) according to the manufacturer’s instructions, by incubating with 1mL of retroviral supernatant (2×10^7^-5×10^7^ TU/mL) and centrifugation at 3200 RPM, 32 °C for two hours. The supernatant was subsequently aspirated off of the wells and 0.5×10^6^ T cells were added in 1mL of T cell media comprised of: AIM V (Thermo Fisher), 5% fetal bovine serum (FBS), 100 U/mL penicillin (Gibco), 100 mg/mL streptomycin (Gibco), 2 mM L-glutamine (Gibco), 10 mM HEPES (Gibco), and 100 U/mL rhIL-2 (Peprotech). After addition of the T cells, the plates were gently spun down at 1200 RPM for 2 min then incubated for 24hrs at 37°C 5% CO_2_. This transduction process was repeated at Day 3 and Day 4 (if necessary). Dynabeads were removed on Day 4 or Day 5 by magnetic separation. Cells were maintained between 0.4 - 2×10^6^ cells/mL and expanded until Day 10. Typically, T cells were transduced with CAR or TCR on Day 2, and then CD47 variants on Days 3 and 4.

### Flow cytometry

Recombinant B7H3-Fc and Her2-Fc (R&D systems) were used to detection B7H3 and Her2 surface CAR, respectively. Likewise, anti-FMC63 and anti-14g2a idiotype antibodies were used to detect CD19 and GD2 CAR, respectively. CAR detection reagents were fluorescently labeled with the DyLight 650 Microscale Antibody Labeling Kit (Thermo Fisher). Anti-FLAG (BioLegend) was used to detect the PIP CAR. NY-ESO-1 TCR was detected with antibodies specific for Vβ13.1 (BioLegend), the beta chain of the NY-ESO-1 TCR. CD47 was detected with B6H12 (BD and Invitrogen), TJC4, Hu5F9, CV-1-Fc, mSIRPɑ-Fc (Sino Biological), or hSIRPɑ-Fc (Sino Biological), followed by detection with an anti-mouse- or human-Fc antibody. The following antibodies were used for detection of cell-surface proteins: calreticulin (FMC 75 mAb; Abcam); human CD4 (clone SK3; BD); human CD8 (clone SK1; BD); human CD45 (clone HI30; Thermo Fisher); mouse CD45 (clone I3/2.3; BD); F4/80 (clone BM8; BioLegend); CD11b (clone M1/70; BD). Surface protein was stained by incubation with 3 μg/mL of detection reagents (or at the concentrations indicated in the figures) for 30 min at 4 °C. Flow cytometry was performed on BD Fortessa and BD Accuri instruments.

### Bioluminescence imaging

Mice were administered either 200μL of 15 μg/mL D-luciferin or a 1:40 dilution of Nano-Glo substrate (Promega, diluted in DPBS) by intraperitoneal injection for firefly luciferase and Antares or nanoluciferase imaging, respectively. Images were acquired on an IVIS or Lago imaging system 4 min after injection for fLuc and 8 min after injection for nLuc/Antares using 30 sec exposures and medium binning. If saturated pixels were detected in the image, an additional image was acquired using the auto-expose setting. Total flux was measured using Living Image (Perkin Elmer) or Aura (Spectral Instruments Imaging) software with a region of interest around the body of each mouse. Only non-saturated images were used for quantification of BLI. Mice were randomized prior to T cell administration to ensure uniform distribution of tumor burden between groups. At the end of the experiment, all images were collected into a single sequence on Aura and set to the same luminescence scale.

### Recombinant protein cloning and production

The gWIZ vector with a BM40 signal peptide was used for protein expression. DNA encoding Hu5F9’s (magrolimab’s) heavy chain with an hIgG1 Fc domain, Hu5F9’s light chain, TJC4’s (lemzoparlimab’s) heavy chain with an hIgG1 Fc domain, and TJC4’s light chain were ordered from Integrated DNA Technologies. Heavy and light chains were individually cloned into AscI/BamHI digested gWIZ vector using Gibson assembly. Plasmids were transfected into Expi293F cells (Thermo Fisher Scientific) in a 1:1 ratio of heavy chain:light chain using ExpiFectamine according to the manufacturer’s instructions. Five days after transfection, supernatant was harvested, adjusted to pH 8.0 and sterile-filtered. Hu5F9 and TJC4 were then purified using recombinant Protein A-Sepharose 4B (Thermo Fisher Scientific) buffer exchanged into PBS and concentrated using Amicon Centrifugal Filters (Millipore Sigma). To assess CD47 binding, cells were stained with Hu5F9 or TJC4 and then stained with labeled anti-human secondary antibody (Invitrogen). B6H12 was acquired from Bio X Cell. CV-1 (ALX-222) was acquired from ALX Oncology. Human SIRPɑ-mFc and mouse SIRPɑ-hFc were acquired from Sino Biologic.

### Animal models

NSG mice (NOD.Cg-Prkdc^scid^ Il2rgt^m1Wjl^/SzJ) were purchased from the Jackson Laboratory and bred in house under Stanford University APLAC-approved protocols. Healthy male and female mice were used for *in vivo* experiments between 6 and 10 weeks old at tumor engraftment and were drug naïve, and not involved in previous procedures. Mice were house in sterile cages in a barrier facility at Stanford University with a 12-hour light/dark cycle. Veterinary Services Center (VSC) staff at Stanford University monitored the mice daily and were euthanized when mice manifested persistent hunched posture, persistent scruffy coat, paralysis, impaired mobility, greater than 20% weight loss, if tumors significantly interfered with normal bodily functions, or if they exceeded limits designated in APLAC-approved protocols. Per recommendation by VSC staff, mice with morbidities were supported with 500μL subcutaneous saline, diet gel (DietGel® 76A, ClearH2O), and wet chow.

### MG63.3 osteosarcoma tumor model

1⨉10^6^ MG63.3 cells in 100μL DPBS were injected into the tibia periostea of six- to ten-week-old NSG male or female mice. Starting fifteen days after tumor implantation and after visual confirmation of tumor formation, mice were treated with 400 μg of B6H12 or PBS three times per week by intraperitoneal injection. On day 21, mice were treated with 10⨉10^6^ GD2.BBζ or B7H3.BBζ CAR T cells or no T cells. Tumor progression was measured with digital calipers twice per week. Mice were euthanized according to the criteria described in the Animal Models section.

### D425 medulloblastoma tumor model

Six- to ten-week-old mice were anesthetized with 3% isoflurane (Minrad International) in an induction chamber. Anesthesia on the stereotactic frame (David Kopf Instruments) was maintained at 2% isoflurane delivered through a nose adaptor. D425 medulloblastoma cells were injected at coordinates 2 mm posterior to lambda on midline and 2 mm deep using a blunt-ended needle (75N, 26s/2″/2, 5 μL; Hamilton Co.). Using a microinjection pump (UMP-3; World Precision Instruments), 0.2⨉10^6^ D425-GL cells were injected in a volume of 3 μL at 30 nL/s. After leaving the needle in place for 1 minute, it was retracted at 3 mm/min. Four days after tumor implantation and after confirmation of tumor formation by bioluminescence, mice were randomized and treated with 10⨉10^6^ B7H3.BBζ CAR^+^ T cells or an equivalent number of Mock T cells intravenously by tail vein injection. Starting on day 4, mice were also treated with 400 μg of B6H12 or PBS three times per week by intraperitoneal injection. Tumor progression was monitored by firefly luciferase BLI.

### Nalm6 leukemia tumor models

Six- to ten-week-old NSG male or female mice were implanted with 1⨉10^6^ Nalm6-GL cells by tail vein injection. CAR specificity, treatment doses and times for the specific model, and antibody doses are indicated in the figure legends. Tumor progression was monitored by firefly luciferase BLI. T cells were quantified by nanoluciferase BLI before and after αCD47 treatment and in the blood by flow cytometry, as indicated. Mice were euthanized according to the criteria described in the Animal Models section.

### T cell depletion model

Six- to ten-week-old NSG male or female mice were implanted with 2⨉10^6^ or 5⨉10^6^ CD19.BBζ- nLuc CAR T cells by tail vein injection (day 0). Mice were then treated twice with B6H12 (250 μg) or PBS by intraperitoneal injection on day 3 and day 5. T cells were quantified by nanoluciferase BLI before (2⨉10^6^ dose: day 2; 5⨉10^6^ dose: day 3) and after (2⨉10^6^ dose: day 9; 5⨉10^6^ dose: day 7) anti-CD47 treatment and in the blood by flow cytometry (2⨉10^6^ dose: day 7; 5⨉10^6^ dose: day 6). Mice were euthanized according to the criteria described in the Animal Models section at the conclusion of the experiment.

### CHLA-255 neuroblastoma metastatic tumor model

Six- to ten-week-old NSG male or female mice were implanted with 1⨉10^6^ CHLA-255-GL cells by tail vein injection. Seven days after tumor implantation and after confirmation of tumor formation by bioluminescence, mice were randomized and treated with 2⨉10^6^ B7H3.BBζ-nLuc CAR T cells with endogenous CD47 knocked out (47_KO_) and over-expressing either CD47 WT (47_WT_) or CD47 Q31P (47_E_) or an equivalent number of mock (non-transduced) T cells intravenously by tail vein injection. Mice were then treated three times with B6H12 (250 μg) or PBS by intraperitoneal injection on day 7, day 9, and day 13. Tumor progression was monitored by firefly luciferase BLI. T cells were quantified by nanoluciferase BLI after αCD47 treatment on day 14 and in the blood by flow cytometry on day 15. Mice were euthanized according to the criteria described in the Animal Models section.

### A375 melanoma tumor model

3⨉10^6^ A375 cells in 100μL DPBS were injected into the flanks of six- to ten-week-old NSG male or female mice. Seven (experiment 1; **Extended Data Fig. 7c**) or fourteen (experiment 2; **Fig. 7d**) days after tumor implantation and after visual confirmation of tumor formation, mice were treated with 1⨉10^6^ NY-ESO-1-Antares TCR T cells with endogenous CD47 knocked-out (47_KO_) and over-expressing CD47 Q31P (47_E_), or an equivalent number of mock-Antares T cells intravenously by tail vein injection. Mice were then treated either: (expt. 1) three times with B6H12 (250 μg) or PBS by intraperitoneal injection on day 9, 11, and 14, or (expt. 2) twice with B6H12 (250 μg) or PBS by intraperitoneal injection on day 15 and 19. Tumor progression was monitored by caliper measurement. T cells in experiment 2 were quantified by nanoluciferase BLI before (day 15) and after (day 20) αCD47 treatment and in the blood by flow cytometry on day 20. Mice were euthanized according to the criteria described in the Animal Models section.

### 143B osteosarcoma tumor model

1⨉10^6^ 143B cells in 100μL DPBS were injected into the tibial periosteum of six- to ten-week-old NSG male or female mice. Five days after tumor implantation and after visual confirmation of tumor formation, mice were treated with 4⨉10^6^ Her2.BBζ-Antares CAR T cells with endogenous CD47 knocked-out (47_KO_) and over-expressing either CD47 WT (47_WT_) or CD47 Q31P (47_E_), or an equivalent number of mock-Antares T cells intravenously by tail vein injection. Mice were then treated twice with B6H12 (250 μg) or PBS by intraperitoneal injection on day 7 and day 11. Tumor progression was monitored by caliper measurement. T cells were quantified by nanoluciferase BLI before (day 7) and after (day 13) αCD47 treatment and in the blood by flow cytometry on day 14. Mice were euthanized according to the criteria described in the Animal Models section.

### CAR T cell GvHD model

Six- to ten-week-old NSG male or female mice were implanted 1⨉10^6^ Nalm6-fLuc cells by tail-vein injection. Mice were then treated with 10⨉10^6^ CD19.BBζ CAR T cells on day 4. Half of the cohort of mice received three doses of B6H12 (250ug) over three days after CAR T administration. Mice were monitored for tumor growth by BLI and signs of GvHD, such as alopecia, dyskeratosis, and weight loss^119^. Spleens and skin were extracted surgically. Skin sections per prepared as slides and stained with H&E in the standard way.

### Isolation of T cells from spleens and tumors

Spleens and tumors were harvested and mechanically dissociated using a gentleMACS dissociator (Miltenyi). Single-cell suspensions were made by passing spleens and tumors through a 70μm cell strainer, depleting red blood cells by ACK lysis (Quality Biological Inc.), and further filtration through flow cytometry filter tubes with 35μm cell strainer caps (Falcon). Single cell suspensions were then frozen in CryoStor buffer in liquid nitrogen, or stained and run directly on flow cytometry, staining for Live/Dead, hCD3, hCD45, hCD4, hCD8, as well as CAR.

### Quantification of T cells and cytokines from blood

Mouse blood was collected from the retro-orbital sinus into Microvette blood collection tubes with EDTA (Fisher Scientific). Red blood cells were depleted by ACK lysis (Quality Biological Inc.), followed by two washes with FACS buffer (PBS + 2% FBS). Samples were stained with anti-hCD45, anti-hCD4, anti-hCD8, anti-hCD47, and anti-CAR reagents. Samples were mixed with CountBright Absolute Counting beads (Thermo Fisher) before flow cytometry analysis. IL-2, IL-6, IL-10, IFNγ, and TNF-ɑ cytokine levels in blood were quantified by LEGENDPlex immunoassays (Biolegend) according to the manufacturer’s instructions. Negative cytokine values were set to 0.

### CRISPR/Cas9 knock-out of CD47

Ribonucleoprotein (RNP) was prepared using synthetic sgRNA with 2′-*O*-methyl phosphorothioate modification (Synthego) diluted in TE buffer at 120 μM. Five microliters sgRNA were incubated with 2.5 μl duplex buffer (IDT) and 2.5 μg Alt-R S.p. Cas9 Nuclease V3 (IDT) for 30 min at room temperature. One hundred–microliter reactions were assembled with 5 million T cells, 90 μl P3 buffer (Lonza), and 10 μl RNP. Cells were pulsed with protocol EO115 using the P3 Primary Cell 4D-Nucleofector Kit and 4D-Nucleofector System (Lonza). Cells were recovered immediately with warm media for 6 hours before transduction with CAR. Guide sequence: CD47-sg3: 5′ AUGCUUUGUUACUAAUAUGG 3′

### Macrophage depletion and peritoneal lavage

Six- to ten-week-old NSG male or female mice were pre-treated by intravenous injection with 200 µL of clodronate liposomes (Liposoma), followed by 400 µg of anti-CSF1R (Bio X Cell, AFS98) by intraperitoneal injection. Mice were treated with 400 µg of anti-CSF1R three times per week for the duration of the experiment. Six days after clodronate treatment, mice were administered with 2⨉10^6^ CD19.28ζ-nLuc CAR T cells, followed by 250 µg B6H12 on day 7. T cells were quantified by nanoluciferase BLI before (day 7) and after (day 9) anti-CD47 treatment. Peritoneal lavage was performed on day 13 with 10 mL of FACS buffer and a 25-gauge needle. Peritoneal cells were collected and stained for Live/Dead, CD11b, F4/80, hCD45, and mCD45, before being run on flow cytometry.

### Incucyte tumor killing assay

5⨉10^4^ GFP-labeled tumor cells were cocultured with 5⨉10^4^ CAR T cells in 200μL RPMI supplemented with 10% FBS, 10mM HEPES, 2mM L-glutamine, 100 U/mL penicillin, and 100μg/mL streptomycin. Triplicate wells were plated in 96-well flat-bottom plates for each condition. Tumor fluorescence was monitored every 2-3 hours with a 10x objective using the Incucyte Zoom system (Essen Bioscience), housed in a cell culture incubator at 37°C and 5% CO_2_, set to take 4 images per well at each time point. Total integrated GFP intensity was quantified using the Zoom software (Essen Bioscience). Data were normalized to the first timepoint and plotted as fold change in tumor fluorescence over time.

### Imaging of patient CSF samples

A cerebrospinal fluid cytospin preparation was collected from a patient treated with axicabtagene ciloleucel (axi-cel) CD19.28ζ CAR T cell therapy, stained with Wright Giemsa, and imaged via microscopy at 1000x magnification, capturing histiocytes with engulfed lymphocytes.

### Single cell analysis of patient samples

Two datasets were re-analyzed: Good, Z., et. al. 2022^48^: scRNA-seq data collected from nine LBCL patients treated with axicabtagene ciloleucel (axi-cel) CD19.28ζ CAR T cell therapy, where 50,000-70,000 CAR T cells (single live CD4^+^ or CD8α^+^ CD235a^−^ CAR^+^ events) were FACS sorted to ≥95% purity and analyzed on the 10x Genomics platform^48, 120^ (GSE168940); and Majzner, R.G., et. al. 2022^49^: scRNA-seq data collected from four DMG patients treated with GD2.BBζ CAR T cell therapy, where cells from the manufacturing product and CSF were analyzed on the 10x Genomics platform^49, 120^ (GSE186802). Where indicated, previously annotated CAR mRNA-expressing cells were used.

### Histology of tissue samples

The tissues assessed include skin and lung. Tissues were harvested and immersion-fixed in 10% neutral buffered formalin. After fixation, tissues were routinely processed, embedded in paraffin, sectioned at 5.0 μm and routinely stained with hematoxylin and eosin (H&E). Tissues were visualized with an Olympus BX43 upright bright-field microscope, and images were captured using an Olympus DP27 camera and cellSens software.

### PIP-CAR toxicity model

PIP-CAR vectors were made as described in the viral vector construction section. Six- to ten-week-old NSG male or female mice were treated with the PIP-CAR T or control CD19.28ζ (non-tumor targeting), Her2.BBζ (tumor targeting), or mock T cells at the dose indicated in the figure legends (10⨉10^6^, 5⨉10^6^, or 2⨉10^6^ CAR T cells) by tail vein injection. Mice in PIP.28ζ and PIP.BBζ groups experienced rapid onset of toxicity (within 1 - 5 days, depending on dose) observed clinically as hunched posture, scruffy coat, slow movement, dehydration, and weight loss. Treatment-related toxicity was monitored by weight change, which was measured prior to T cell administration and 1-2⨉ per week thereafter. % weight change was calculated according to: % weight change = (weight at time x) / (initial weight)−1) ⨉ 100. Mice died from toxicity or were euthanized if they reached 20% weight loss or showed clinical signs of severe toxicity, as described in the Animal Models section.

For assessment of T cell localization and activation, CD19.28ζ-nLuc or PIP.28ζ-nLuc CAR T cells were transduced with a firefly luciferase reporter under control of an NFκB inducible promoter. Mice were implanted with 5⨉10^6^ CAR-T cells and imaged daily with Nano-GLO substrate (nLuc; total CAR T) and luciferin (fLuc; active CAR T), with each substrate dose separated by twelve hours. For organ BLI analysis, four days after treatment with 5⨉10^6^ CD19.28ζ-nLuc or PIP.28ζ-nLuc, mice were injected with either Nano-GLO substrate (nLuc; total CAR T) or luciferin (fLuc; active CAR T) and euthanized 10 minutes after injection. Organs were harvested following standard procedure and imaged using BLI on a IVIS machine (Perkin Elmer) in 6-well plates. For safety-switch models, mice dosed with 2⨉10^6^ PIP.28ζ CAR T cells were administered 250 µg B6H12 over three consecutive days (days 2, 3, and 4), two days after CAR administrations (day 0) as indicated in the figure. For tumor-bearing PIP toxicity models, 0.5⨉10^6^ 143B osteosarcoma cells were engrafted in the tibia periosteum of NSG mice five days prior to T cell administration. Tumor progression was monitored by caliper measurements.

### Yeast surface display vectors

A DNA sequence encoding the CD47 Ig-like domain (Gln19 – Ser135) was cloned into the pCTCON2 yeast-surface display vector (Addgene) using the NheI and BamHI sites. The pFreeNTerm (pFNT) vector was based on the pCL backbone^121^, designing an intrinsic NheI cutsite into the Aga2p signal sequence as the 5’ cloning site and using a MluI cutsite prior to a Gly_4_Ser 3⨉ linker as the 3’ cloning site. The CD47 Ig-like domain (Gln19 – Ser135) was cloned into the pFNT yeast-surface display vector using these NheI and MluI sites.

### Yeast surface display binding assays

EBY100 yeast were transformed with pCTCON2 or pFNT plasmids and selected on SD-CAA-Agar plates. Yeast (∼100,000 per sample) were grown and induced in SG-CAA, and binding set up over a range of soluble ligand or receptor concentrations in phosphate-buffered saline (PBS) containing 1 mg ml^−1^ bovine serum albumin (BSA; BPBS), taking into account ligand depletion and equilibrium time^122^. After incubation with binding partner, yeast cells were washed once with BPBS, then incubated with a 1:5,000 dilution of chicken anti-c-myc antibody (A21281, Invitrogen) for pCTCON2 displayed proteins, and incubated for 30 min at 4°C in the dark. After primary addition, samples were washed once with BPBS, and secondary antibodies were added. Expression was detected with a 1:500 dilution of goat anti-chicken Alexa Fluor 488 or Alexa Fluor 647 (Invitrogen). For pFNT displayed proteins, co-displayed GFP was used to monitor expression. Binding of proteins with mouse Fc domains (hSIRPɑ, B6H12) was detected with a 1:500 dilution of goat anti-mouse Alexa Fluor 488 or Alexa Fluor 647 (Invitrogen). Binding of proteins with a human Fc domain (CV-1, mSIRPɑ, Hu5F9, TJC4) was detected using a 1:500 dilution of goat anti-human Alexa Fluor 488 or Alexa Fluor 647 (Invitrogen). Secondary antibodies were incubated for 15 min at 4°C in the dark. After secondary incubation, samples were washed once with BPBS, pelleted, and left pelleted on ice until analysis. Samples were analyzed by resuspending them in 50 μL of BPBS and running flow cytometry using a BD Accuri C6 (BD Biosciences). Samples were gated for bulk yeast cells (forward scatter (FSC) vs. side scatter (SSC)) and then for single cells (FSC-Height vs. FSC-Area). Expressing yeast were determined and gated via C-terminal c-myc tag or GFP detection. The geometric mean of the binding fluorescence signal was quantified from the expressing population and used as a raw binding value. When comparing binding signals, the average fluorescence expression signal was quantified for different protein variants and used to normalize binding signal. To determine “fraction bound,” binding signals were divided by the signal derived from the highest concentration of binding partner used, or that derived from binding to wild-type CD47. To calculate K_d_ values, data were analyzed in GraphPad Prism (v9.3.1) using non-linear regression curve fit.

### Yeast surface display library generation, sorting, and sequencing

CD47 was expressed in *Saccharomyces cerevisiae* (strain: EBY100; ATCC MYA-4941) as a genetic fusion to the agglutinin mating protein Aga2p. An error-prone PCR library was created using the CD47 Ig-like domain (Gln19 to Ser135) as a template and mutations were introduced with a Gene Morph II random mutagenesis kit (Aglilent), following the manufacturer’s instructions. Separate PCRs were performed using various concentrations of Mutazyme II enzyme. Products from these reactions were purified via gel electrophoresis, pooled, and amplified with standard PCR using Phusion polymerase (New England BioLabs). Purified mutant DNA and linearized plasmid were electroporated into EBY100 yeast, where they were assembled *in vivo* through homologous recombination. We estimated 5×10^7^ variants for the library, determined by dilution plating and colony counting. Yeast were grown in SD-CAA media and induced for CD47 protein expression by growth in media containing 90% SG-CAA and 10% SD-CAA overnight^122^. Yeast displaying CD47 variants were isolated via fluorescence-activated cell sorting (FACS) using a SONY SH800S cell sorter (SONY) and analyzed with a BD Accuri C6 flow cytometer (BD Biosciences). Data were analyzed using FlowJo software (v 10.6.1, Tree Star Inc.). Screens were carried out using equilibrium binding conditions where yeast were incubated at room temperature in BPBS with the following concentrations of B6H12 or CV-1 for two hours. For negative sorts to B6H12, the CD47-expressing, but non-binding populations of yeast were collected. For positive sorts to CV-1, the CD47-expressing and binding populations of yeast were collected. Sort 1, negative sort, 500 pM B6H12; Sort 2, negative sort, 5 nM B6H12; Sort 3, positive sort, 20 nM CV-1; Sort 4, negative sort, 20 nM B6H12; Sort 5, negative sort, 50 nM B6H12; Sort 6, positive sort, 10 nM CV-1. After incubation with B6H12 or CV-1, yeast were pelleted, washed, and labeled with fluorescent antibodies as described above prior to sorting. Sorted yeast clones were propagated, induced for CD47 expression, and subjected to iterative rounds of FACS as described above. After each round of screening, plasmid DNA was recovered using a Zymoprep yeast plasmid miniprep I kit (Zymo Research Corp), transformed into DH10B electrocompetent cells (Thermo Fisher), and isolated using a GeneJET plasmid miniprep kit (Thermo Fisher). Sequencing was performed by ELIM Biopharmaceuticals, Inc. (Hayward, CA).

### CD47 structure modeling

CD47 structures were downloaded from the protein data bank (PDB) and analyzed using PyMol. The CD47-hSIRPa structure used was 2JJS^58^. The CD47-B6H12 structure used was 5TZU^59^.

### Phagocytosis assay

For all flow-based *in vitro* phagocytosis assays, T cells and human macrophages were co-cultured at a ratio of 2:1 (e.g. 100,000 T cells : 50,000 macrophages) in ultra-low-attachment 96-well U-bottom plates (Corning) in serum-free RPMI (Thermo Fisher Scientific). T cells were labeled with CFSE (Invitrogen) by suspending cells in PBS (5 µM working solution) as per manufacturer instructions for 20 min at 37 °C protected from light and washed twice with 20 ml of FBS-containing media before co-culture. Cells were then either incubated alone or in the presence of anti-CD47 (clone B6H12, Bio X Cell) at a concentration of 10 μg ml^−1^. T cells and antibodies were incubated for 30 min in a humidified 5% CO_2_ incubator at 37 °C. Plates were washed two times; human macrophages were added to the plate; and plates were incubated for 1–2 h at 37 °C. Phagocytosis was stopped by washing with 4 °C PBS and centrifugation at 336g before the cells were stained with Live/Dead stain and anti-CD11b-APC. Assays were analyzed by flow cytometry, and phagocytosis was measured as the number of CD11b^+^ and CFSE^+^ macrophages, quantified as a percentage of the total CD11b^+^ macrophages and normalized to the control condition.

### 143B correlative study and tumor dissociation

1⨉10^6^ 143B cells in 100μL DPBS were injected into the tibia periosteum of six-to ten-week-old NSG mice. Thirteen days after tumor implantation and after visual confirmation of tumor formation, mice were treated with 4⨉10^6^ Her2.BBζ CAR-T cells with endogenous CD47 knocked-out (47_KO_) and over-expressing either CD47 WT (47_WT_) or CD47 Q31P (47_E_), an equivalent number of Mock T cells intravenously by tail vein injection, or no T cells. Mice were then treated twice with B6H12 (250 µg) or PBS by intraperitoneal injection on day 15 and day 19. Tumor progression was monitored by caliper measurement. Tumors were harvested at day 21 post tumor implantation (day 8 post CAR T treatment). Tumors were weighed and then split with a razor, with one section being fixed in 10% paraformaldehyde, and the other mechanically dissociated using a gentleMACS dissociator (Miltenyi). Single-cell suspensions were made by passing tumors through a 70μm cell strainer, depleting red blood cells by ACK lysis (Quality Biological Inc.), and further filtration through flow cytometry filter tubes with 35 μm cell strainer caps (Falcon). Single cell suspensions were subsequently stained for flow cytometry and FACS. Formaldehyde fixed tumor had paraformaldehyde removed after 24 h and replaced with 70% ethanol for long term storage. Tumor sections were then formalin-fixed and paraffin-embedded following the standard protocol.

### Flow cytometry and IHC on dissociated tumors

Flow cytometry: Tumors were harvested as above. Single cell suspensions of dissociated tumors were stained for CAR (Her2-Fc; R&D), hCD19 (BD), CD11b (BD), F4/80 (BioLegend), hCD45 (Invitrogen), mCD45 (BD), hCD47 (BD), and Live/Dead (Invitrogen) for 30 minutes before being analyzed by flow cytometry.

IHC: Tumors were harvested as above. Formalin-fixed, paraffin-embedded xenograft tumor sections were used. F4/80 (Cell Signaling Technology) staining was performed manually, and hCD3 (Abcam) and Arg1 (Cell Signaling Technology) staining was performed using the Ventana Discovery platform. In brief, tissue sections were incubated in either 6mM citrate buffer (F4/80) or Tris EDTA buffer (CD3/Arg1, 1:100 and 1:250 dilution respectively) (cell conditioning 1 standard) at 100 °C for 25min (F480) or 95 °C for 1h (CD3/Arg1) to retrieve antigenicity, followed by incubation with the respective primary antibody for 1 h. Bound primary antibodies were incubated with the respective secondary antibodies (Vector Laboratories or Jackson Laboratory) with 1:500 dilution, followed by UltraMap HRP and Vectore Lab (F4/80) or ChromoMap DAB (CD3/Arg1) detection. For IHC analysis, tumor regions were identified based on histology. F4/80, CD3, and Arg1 positivity were analyzed for each tumor region. F4/80, CD3, and Arg1 IHC positivity scores were automatically quantified in the regions of interest with Aperio ImageScope software. Regions of interest were randomly selected within the tumor to exclude macrophages present in the normal tissue around the tumor.

### Single cell analysis of dissociated 143B tumors

Dissociated tumors from the 143B osteosarcoma model described above were sorted for live cells using a Live/Dead stain (Invitrogen) at the Stanford Shared FACS facility. Single-cell RNAseq libraries were prepared using the Chromium Next GEM Single Cell 5’ v2 platform (10x GENOMICS). Libraries were sent to Novogene for sequencing on a NovoSeq S4 lane (PE150) with approximately 30,000 mean reads per cell. Reads were aligned and quantified with Cell Ranger (10x GENOMICS) using the standard workflow, with the reference transcriptomes GRCh38 for human and mm10 for mouse. The Cell Ranger output was imported into R using Seurat 4.2.0. The following filters were applied using the subset function to select for live cells: nFeature_RNA > 200 & nFeature_RNA < 5000; percent mitochondrial reads < 5%. After filtering, the eight biological samples ranged from 7658 – 9327 mean unique molecular identifiers (UMI) per cell. The data matrix was normalized with NormalizeData and scaled with Seurat. Differential expression analysis, clustering, and UMAP dimensionality reduction analysis were performed on the resulting data matrix using Seurat^123^. Pathway analysis was performed using Enrichr^124^.

### Statistical Analyses

The specific statistical tests utilized are indicated in the figure legends. Statistical analyses were performed using Prism (v 9.3.1, GraphPad Software). For comparisons between two groups, statistical significance was assayed by two-tailed unpaired Student’s *t*-test or a Mann-Whitney test. For comparison within in vivo studies and between grouped studies, a two-way analysis of variance (ANOVA) combined with Tukey’s multiple comparison test for post hoc analysis was performed. Significance for survival data was calculated using the log-rank Mantel–Cox test. Sample sizes were determined on the basis of the variability of tumor models used. Tumor-bearing animals were assigned to the treatment groups to ensure an equal distribution of tumor sizes between groups. Data are represented as mean ± standard deviation (*in vitro* studies) or mean ± standard error of the mean (some *in vivo* studies). For all statistical analyses, *P* values are indicated in each figure panel.

### Data availability

All data associated with this paper are included in the manuscript and the supplementary materials. scRNA-seq data will be deposited to the Gene Expression Omnibus upon publication of the manuscript.

## References

1 DeNardo, D. G. & Ruffell, B. Macrophages as regulators of tumour immunity and immunotherapy. Nat Rev Immunol 19, 369–382 (2019). https://doi.org:10.1038/s41577-019-0127-6

2 Barry, S. T., Gabrilovich, D. I., Sansom, O. J., Campbell, A. D. & Morton, J. P. Therapeutic targeting of tumour myeloid cells. Nat Rev Cancer 23, 216–237 (2023). https://doi.org:10.1038/s41568-022-00546-2

3 Kloosterman, D. J. & Akkari, L. Macrophages at the interface of the co-evolving cancer ecosystem. Cell 186 (2023). https://doi.org:10.1016/j.cell.2023.02.020

4 Shen, H. et al. Prognostic Value of Tumor-Associated Macrophages in Clear Cell Renal Cell Carcinoma: A Systematic Review and Meta-Analysis. Front Oncol 11, 657318 (2021). https://doi.org:10.3389/fonc.2021.657318

5 Li, J. et al. Tumor-associated macrophage infiltration and prognosis in colorectal cancer: systematic review and meta-analysis. Int J Colorectal Dis 35, 1203–1210 (2020). https://doi.org:10.1007/s00384-020-03593-z

6 Ries, C. H. et al. Targeting tumor-associated macrophages with anti-CSF-1R antibody reveals a strategy for cancer therapy. Cancer Cell 25, 846–859 (2014). https://doi.org:10.1016/j.ccr.2014.05.016

7 Zhu, Y. et al. CSF1/CSF1R blockade reprograms tumor-infiltrating macrophages and improves response to T-cell checkpoint immunotherapy in pancreatic cancer models. Cancer Res 74, 5057–5069 (2014). https://doi.org:10.1158/0008-5472.CAN-13-3723

8 Xu, J. et al. CSF1R signaling blockade stanches tumor-infiltrating myeloid cells and improves the efficacy of radiotherapy in prostate cancer. Cancer Res 73, 2782–2794 (2013). https://doi.org:10.1158/0008-5472.CAN-12-3981

9 Noel, M. et al. Phase 1b study of a small molecule antagonist of human chemokine (C-C motif) receptor 2 (PF-04136309) in combination with nab-paclitaxel/gemcitabine in first-line treatment of metastatic pancreatic ductal adenocarcinoma. Invest New Drugs 38, 800–811 (2020). https://doi.org:10.1007/s10637-019-00830-3

10 Chen, Y. L. Prognostic significance of tumor-associated macrophages in patients with nasopharyngeal carcinoma: A meta-analysis. Medicine (Baltimore) 99, e21999 (2020). https://doi.org:10.1097/MD.0000000000021999

11 Matlung, H. L., Szilagyi, K., Barclay, N. A. & van den Berg, T. K. The CD47-SIRPalpha signaling axis as an innate immune checkpoint in cancer. Immunol Rev 276, 145–164 (2017). https://doi.org:10.1111/imr.12527

12 Majeti, R. et al. CD47 is an adverse prognostic factor and therapeutic antibody target on human acute myeloid leukemia stem cells. Cell 138, 286–299 (2009). https://doi.org:10.1016/j.cell.2009.05.045

13 Chao, M. P. et al. Anti-CD47 antibody synergizes with rituximab to promote phagocytosis and eradicate non-Hodgkin lymphoma. Cell 142, 699–713 (2010). https://doi.org:10.1016/j.cell.2010.07.044

14 Willingham, S. B. et al. The CD47-signal regulatory protein alpha (SIRPa) interaction is a therapeutic target for human solid tumors. Proceedings of the National Academy of Sciences of the United States of America 109, 6662–6667 (2012). https://doi.org:10.1073/PNAS.1121623109/-/DCSUPPLEMENTAL

15 Advani, R. et al. CD47 Blockade by Hu5F9-G4 and Rituximab in Non-Hodgkin’s Lymphoma. New England Journal of Medicine 379, 1711–1721 (2018). https://doi.org:10.1056/NEJMOA1807315/SUPPL_FILE/NEJMOA1807315_DISCLOSURES.PDF

16 Sallman, D. A. et al. Magrolimab in Combination With Azacitidine in Patients With Higher-Risk Myelodysplastic Syndromes: Final Results of a Phase Ib Study. J Clin Oncol, JCO2201794 (2023). https://doi.org:10.1200/JCO.22.01794

17 Sikic, B. I. et al. First-in-Human, First-in-Class Phase I Trial of the Anti-CD47 Antibody Hu5F9-G4 in Patients With Advanced Cancers. J Clin Oncol 37, 946–953 (2019). https://doi.org:10.1200/JCO.18.02018

18 Lakhani, N. J. et al. Evorpacept alone and in combination with pembrolizumab or trastuzumab in patients with advanced solid tumours (ASPEN-01): a first-in-human, open-label, multicentre, phase 1 dose-escalation and dose-expansion study. Lancet Oncol 22, 1740–1751 (2021). https://doi.org:10.1016/S1470-2045(21)00584-2

19 Neelapu, S. S. et al. Axicabtagene Ciloleucel CAR T-Cell Therapy in Refractory Large B-Cell Lymphoma. N Engl J Med 377, 2531–2544 (2017). https://doi.org:10.1056/NEJMoa1707447

20 Fry, T. J. et al. CD22-targeted CAR T cells induce remission in B-ALL that is naive or resistant to CD19-targeted CAR immunotherapy. Nat Med 24, 20–28 (2018). https://doi.org:10.1038/nm.4441

21 Maude, S. L. et al. Tisagenlecleucel in Children and Young Adults with B-Cell Lymphoblastic Leukemia. N Engl J Med 378, 439–448 (2018). https://doi.org:10.1056/NEJMoa1709866

22 Abramson, J. S. et al. Lisocabtagene maraleucel for patients with relapsed or refractory large B-cell lymphomas (TRANSCEND NHL 001): a multicentre seamless design study. Lancet 396, 839–852 (2020). https://doi.org:10.1016/S0140-6736(20)31366-0

23 Wang, M. et al. KTE-X19 CAR T-Cell Therapy in Relapsed or Refractory Mantle-Cell Lymphoma. N Engl J Med 382, 1331–1342 (2020). https://doi.org:10.1056/NEJMoa1914347

24 Berdeja, J. G. et al. Ciltacabtagene autoleucel, a B-cell maturation antigen-directed chimeric antigen receptor T-cell therapy in patients with relapsed or refractory multiple myeloma (CARTITUDE-1): a phase 1b/2 open-label study. Lancet 398, 314–324 (2021). https://doi.org:10.1016/S0140-6736(21)00933-8

25 Munshi, N. C. et al. Idecabtagene Vicleucel in Relapsed and Refractory Multiple Myeloma. N Engl J Med 384, 705–716 (2021). https://doi.org:10.1056/NEJMoa2024850

26 Fowler, N. H. et al. Tisagenlecleucel in adult relapsed or refractory follicular lymphoma: the phase 2 ELARA trial. Nat Med 28, 325–332 (2022). https://doi.org:10.1038/s41591-021-01622-0

27 Kamdar, M. et al. Lisocabtagene maraleucel versus standard of care with salvage chemotherapy followed by autologous stem cell transplantation as second-line treatment in patients with relapsed or refractory large B-cell lymphoma (TRANSFORM): results from an interim analysis of an open-label, randomised, phase 3 trial. Lancet 399, 2294–2308 (2022). https://doi.org:10.1016/S0140-6736(22)00662-6

28 Locke, F. L. et al. Axicabtagene Ciloleucel as Second-Line Therapy for Large B-Cell Lymphoma. N Engl J Med 386, 640–654 (2022). https://doi.org:10.1056/NEJMoa2116133

29 Nastoupil, L. J. et al. Standard-of-Care Axicabtagene Ciloleucel for Relapsed or Refractory Large B-Cell Lymphoma: Results From the US Lymphoma CAR T Consortium. J Clin Oncol 38, 3119–3128 (2020). https://doi.org:10.1200/JCO.19.02104

30 Schultz, L. M. et al. Disease Burden Affects Outcomes in Pediatric and Young Adult B-Cell Lymphoblastic Leukemia After Commercial Tisagenlecleucel: A Pediatric Real-World Chimeric Antigen Receptor Consortium Report. J Clin Oncol 40, 945–955 (2022). https://doi.org:10.1200/JCO.20.03585

31 Labanieh, L. & Mackall, C. L. CAR immune cells: design principles, resistance and the next generation. Nature 2023 614:7949 614, 635–648 (2023). https://doi.org:10.1038/s41586-023-05707-3

32 Siegel, R. L., Miller, K. D. & Jemal, A. Cancer statistics, 2020. CA Cancer J Clin 70, 7–30 (2020). https://doi.org:10.3322/caac.21590

33 Schmidts, A. & Maus, M. V. Making CAR T Cells a Solid Option for Solid Tumors. Front Immunol 9, 2593 (2018). https://doi.org:10.3389/fimmu.2018.02593

34 Majzner, R. G. et al. CAR T cells targeting B7-H3, a pan-cancer antigen, demonstrate potent preclinical activity against pediatric solid tumors and brain tumors. Clinical Cancer Research 25, 2560–2574 (2019). https://doi.org:10.1158/1078-0432.CCR-18-0432/73008/AM/CAR-T-CELLS-TARGETING-B7-H3-A-PAN-CANCER-ANTIGEN

35 Labanieh, L. et al. Enhanced safety and efficacy of protease-regulated CAR-T cell receptors. Cell 185, 1745–1763.e1722 (2022). https://doi.org:10.1016/J.CELL.2022.03.041

36 Theruvath, J. et al. Anti-GD2 synergizes with CD47 blockade to mediate tumor eradication. Nature Medicine 2022 28:2 28, 333–344 (2022). https://doi.org:10.1038/s41591-021-01625-x

37 Gresham, H. D., Goodwin, J. L., Allen, P. M., Anderson, D. C. & Brown, E. J. A novel member of the integrin receptor family mediates Arg-Gly-Asp-stimulated neutrophil phagocytosis. Journal of Cell Biology 108, 1935–1943 (1989). https://doi.org:10.1083/JCB.108.5.1935

38 Lo, J. et al. Anti-CD47 antibody suppresses tumour growth and augments the effect of chemotherapy treatment in hepatocellular carcinoma. Liver International 36, 737–745 (2016). https://doi.org:10.1111/LIV.12963/SUPPINFO

39 Kaur, S. et al. A function-blocking CD47 antibody suppresses stem cell and EGF signaling in triple-negative breast cancer. Oncotarget 7, 10133–10152 (2016). https://doi.org:10.18632/ONCOTARGET.7100

40 Thomas, R. et al. NY-ESO-1 Based Immunotherapy of Cancer: Current Perspectives. Frontiers in Immunology 9, 947–947 (2018). https://doi.org:10.3389/FIMMU.2018.00947

41 Chu, J. et al. A bright cyan-excitable orange fluorescent protein facilitates dual-emission microscopy and enhances bioluminescence imaging in vivo. Nature Biotechnology 2016 34:7 34, 760–767 (2016). https://doi.org:10.1038/nbt.3550

42 Musolino, A. et al. Role of Fcγ receptors in HER2-targeted breast cancer therapy. (2022). https://doi.org:10.1136/jitc-2021-003171

43 Lo, M. et al. Effector-attenuating Substitutions That Maintain Antibody Stability and Reduce Toxicity in Mice. The Journal of Biological Chemistry 292, 3900–3900 (2017). https://doi.org:10.1074/JBC.M116.767749

44 Weiskopf, K. et al. Engineered SIRPα variants as immunotherapeutic adjuvants to anticancer antibodies. Science 341, 88–91 (2013). https://doi.org:10.1126/SCIENCE.1238856/SUPPL_FILE/WEISKOPF.SM.PDF

45 Hu, X. et al. Engineered Hypoimmune Allogeneic CAR T Cells Exhibit Innate and Adaptive Immune Evasion Even after Sensitization in Humanized Mice and Retain Potent Anti-Tumor Activity. Blood 138, 1690–1690 (2021). https://doi.org:10.1182/BLOOD-2021-150021

46 Jaiswal, S. et al. CD47 Is Upregulated on Circulating Hematopoietic Stem Cells and Leukemia Cells to Avoid Phagocytosis. Cell 138, 271–285 (2009). https://doi.org:10.1016/J.CELL.2009.05.046/ATTACHMENT/C76B88E8-D0AF-494A-9683-993A7A88A224/MMC1.PDF

47 Chao, M. P. et al. Calreticulin is the dominant pro-phagocytic signal on multiple human cancers and is counterbalanced by CD47. Science translational medicine 2, 63ra94-63ra94 (2010). https://doi.org:10.1126/SCITRANSLMED.3001375

48 Good, Z. et al. Post-infusion CAR TReg cells identify patients resistant to CD19-CAR therapy. Nature Medicine 2022 28:9 28, 1860–1871 (2022). https://doi.org:10.1038/s41591-022-01960-7

49 Majzner, R. G. et al. GD2-CAR T cell therapy for H3K27M-mutated diffuse midline gliomas. Nature 603, 934–941 (2022). https://doi.org:doi:10.1038/s41586-022-04489-4

50 Miller, C. L. et al. Systemic delivery of a targeted synthetic immunostimulant transforms the immune landscape for effective tumor regression. Cell Chemical Biology 29, 451–462.e458 (2022). https://doi.org:10.1016/J.CHEMBIOL.2021.10.012

51 Kimura, R. H., Levin, A. M., Cochran, F. V. & Cochran, J. R. Engineered cystine knot peptides that bind αvβ3, αvβ5, and α5β1 integrins with low-nanomolar affinity. Proteins: Structure, Function, and Bioinformatics 77, 359–369 (2009). https://doi.org:10.1002/PROT.22441

52 Van Agthoven, J. F. et al. Structural Basis of the Differential Binding of Engineered Knottins to Integrins αVβ3 and α5β1. Structure 27, 1443–1451.e1446 (2019). https://doi.org:10.1016/J.STR.2019.06.011

53 Moore, S. J. et al. Engineered knottin peptide enables noninvasive optical imaging of intracranial medulloblastoma. Proceedings of the National Academy of Sciences 110, 14598–14603 (2013). https://doi.org:10.1073/PNAS.1311333110

54 Ali, N. et al. Xenogeneic Graft-versus-Host-Disease in NOD-scid IL-2Rγnull Mice Display a T-Effector Memory Phenotype. PLOS ONE 7, e44219–e44219 (2012). https://doi.org:10.1371/JOURNAL.PONE.0044219

55 Kamiya, T., Wong, D., Png, Y. T. & Campana, D. A novel method to generate T-cell receptor–deficient chimeric antigen receptor T cells. Blood Advances 2, 517–528 (2018). https://doi.org:10.1182/BLOODADVANCES.2017012823

56 Liu, P. et al. Acute Graft-Versus-Host Disease After Humanized Anti-CD19-CAR T Therapy in Relapsed B-ALL Patients After Allogeneic Hematopoietic Stem Cell Transplant. Frontiers in Oncology 10, 1975–1975 (2020). https://doi.org:10.3389/FONC.2020.573822/BIBTEX

57 Ho, C. C. M. et al. “Velcro” Engineering of High Affinity CD47 Ectodomain as Signal Regulatory Protein α (SIRPα) Antagonists That Enhance Antibody-dependent Cellular Phagocytosis *. Journal of Biological Chemistry 290, 12650–12663 (2015). https://doi.org:10.1074/JBC.M115.648220

58 Hatherley, D. et al. Paired receptor specificity explained by structures of signal regulatory proteins alone and complexed with CD47. Molecular cell 31, 266–277 (2008). https://doi.org:10.1016/J.MOLCEL.2008.05.026

59 Pietsch, E. C. et al. Anti-leukemic activity and tolerability of anti-human CD47 monoclonal antibodies. Blood Cancer Journal 7, e536–e536 (2017). https://doi.org:10.1038/BCJ.2017.7

60 Fenalti, G. et al. Structure of the human marker of self 5-transmembrane receptor CD47. Nature communications 12 (2021). https://doi.org:10.1038/S41467-021-25475-W

61 Mehta, A. et al. Lemzoparlimab, a Differentiated Anti-CD47 Antibody in Combination with Rituximab in Relapsed and Refractory Non-Hodgkin’s Lymphoma: Initial Clinical Results. Blood 138, 3542–3542 (2021). https://doi.org:10.1182/BLOOD-2021-150606

62 Li, M. et al. Anti-CD47 immunotherapy in combination with BCL-2 inhibitor to enhance anti-tumor activity in B-cell lymphoma. Hematological Oncology 40, 596–608 (2022). https://doi.org:10.1002/HON.3009

63 Sikic, B. I. et al. First-in-Human, First-in-Class Phase I Trial of the Anti-CD47 Antibody Hu5F9-G4 in Patients With Advanced Cancers. Journal of Clinical Oncology 37, 946–946 (2019). https://doi.org:10.1200/JCO.18.02018

64 Weiskopf, K. et al. CD47-blocking immunotherapies stimulate macrophage-mediated destruction of small-cell lung cancer. The Journal of Clinical Investigation 126, 2610–2610 (2016). https://doi.org:10.1172/JCI81603

65 Meng, Z., Wang, Z., Guo, B., Cao, W. & Shen, H. TJC4, a Differentiated Anti-CD47 Antibody with Novel Epitope and RBC Sparing Properties. Blood 134, 4063–4063 (2019). https://doi.org:10.1182/BLOOD-2019-122793

66 Appay, V. & Rowland-Jones, S. L. RANTES: a versatile and controversial chemokine. Trends in Immunology 22, 83–87 (2001). https://doi.org:10.1016/S1471-4906(00)01812-3

67 Menten, P., Wuyts, A. & Van Damme, J. Macrophage inflammatory protein-1. Cytokine & Growth Factor Reviews 13, 455–481 (2002). https://doi.org:10.1016/S1359-6101(02)00045-X

68 Kersten, K. et al. Spatiotemporal co-dependency between macrophages and exhausted CD8+ T cells in cancer. Cancer Cell 40, 624–638.e629 (2022). https://doi.org:10.1016/J.CCELL.2022.05.004

69 Fitzgerald, K. A., O’Neill, L. A. J. & Gearing, A. J. H. The Cytokine Factsbook and Webfacts (2nd Edition). 526–526 (2001).

70 Zou, J. J. et al. Structure-Function Analysis of the p35 Subunit of Mouse Interleukin 12. Journal of Biological Chemistry 270, 5864–5871 (1995). https://doi.org:10.1074/jbc.270.11.5864

71 Grewal, I. S. & Flavell, R. A. The role of CD40 ligand in costimulation and T-cell activation. Immunological reviews 153 (1996). https://doi.org:10.1111/j.1600-065x.1996.tb00921.x

72 Kuhn, N. F. et al. CD40 Ligand-Modified Chimeric Antigen Receptor T Cells Enhance Antitumor Function by Eliciting an Endogenous Antitumor Response. Cancer Cell 35, 473–488.e476 (2019). https://doi.org:10.1016/j.ccell.2019.02.006

73 Duan, Z. & Luo, Y. Targeting macrophages in cancer immunotherapy. Signal Transduction and Targeted Therapy 2021 6:1 6, 1–21 (2021). https://doi.org:10.1038/s41392-021-00506-6

74 Zhang, Z. et al. Role of lysosomes in physiological activities, diseases, and therapy. Journal of Hematology & Oncology 14, 1–39 (2021). https://doi.org:doi:10.1186/s13045-021-01087-1

75 Yao, Y., Xu, X. H. & Jin, L. Macrophage polarization in physiological and pathological pregnancy. Frontiers in Immunology 10, 792–792 (2019). https://doi.org:10.3389/FIMMU.2019.00792/BIBTEX

76 Zizzo, G., Hilliard, B. A., Monestier, M. & Cohen, P. L. Efficient Clearance of Early Apoptotic Cells by Human Macrophages Requires M2c Polarization and MerTK Induction. The Journal of Immunology 189, 3508–3520 (2012). https://doi.org:10.4049/JIMMUNOL.1200662

77 Roszer, T. Understanding the mysterious M2 macrophage through activation markers and effector mechanisms. Mediators of Inflammation 2015 (2015). https://doi.org:10.1155/2015/816460

78 Blazar, B. R. et al. CD47 (integrin-associated protein) engagement of dendritic cell and macrophage counterreceptors is required to prevent the clearance of donor lymphohematopoietic cells. J Exp Med 194, 541–549 (2001). https://doi.org:10.1084/jem.194.4.541

79 Deuse, T. et al. Hypoimmunogenic derivatives of induced pluripotent stem cells evade immune rejection in fully immunocompetent allogeneic recipients. Nat Biotechnol 37, 252–258 (2019). https://doi.org:10.1038/s41587-019-0016-3

80 Hu, X. et al. Human hypoimmune primary pancreatic islets avoid rejection and autoimmunity and alleviate diabetes in allogeneic humanized mice. Sci Transl Med 15, eadg5794 (2023). https://doi.org:10.1126/scitranslmed.adg5794

81 Hu, X. et al. Hypoimmune anti-CD19 chimeric antigen receptor T cells provide lasting tumor control in fully immunocompetent allogeneic humanized mice. Nat Commun 14, 2020 (2023). https://doi.org:10.1038/s41467-023-37785-2

82 Beckett, A. N. et al. CD47 expression is critical for CAR T-cell survival in vivo. J Immunother Cancer 11 (2023). https://doi.org:10.1136/jitc-2022-005857

83 Li, L. X., Atif, S. M., Schmiel, S. E., Lee, S. J. & McSorley, S. J. Increased susceptibility to Salmonella infection in signal regulatory protein alpha-deficient mice. J Immunol 189, 2537–2544 (2012). https://doi.org:10.4049/jimmunol.1200429

84 Barclay, A. N. & Van den Berg, T. K. The interaction between signal regulatory protein alpha (SIRPalpha) and CD47: structure, function, and therapeutic target. Annu Rev Immunol 32, 25–50 (2014). https://doi.org:10.1146/annurev-immunol-032713-120142

85 Autio, A. et al. SIRPalpha - CD47 axis regulates dendritic cell-T cell interactions and TCR activation during T cell priming in spleen. PLoS One 17, e0266566 (2022). https://doi.org:10.1371/journal.pone.0266566

86 Peluso, M. O. et al. The Fully human anti-CD47 antibody SRF231 exerts dual-mechanism antitumor activity via engagement of the activating receptor CD32a. J Immunother Cancer 8 (2020). https://doi.org:10.1136/jitc-2019-000413

87 Goswami, S., Anandhan, S., Raychaudhuri, D. & Sharma, P. Myeloid cell-targeted therapies for solid tumours. Nat Rev Immunol 23, 106–120 (2023). https://doi.org:10.1038/s41577-022-00737-w

88 Hong, D. S. et al. Eganelisib, a First-in-Class PI3K-gamma Inhibitor, in Patients with Advanced Solid Tumors: Results of the Phase 1/1b MARIO-1 Trial. Clin Cancer Res (2023). https://doi.org:10.1158/1078-0432.CCR-22-3313

89 Gholamin, S. et al. Disrupting the CD47-SIRPalpha anti-phagocytic axis by a humanized anti-CD47 antibody is an efficacious treatment for malignant pediatric brain tumors. Sci Transl Med 9 (2017). https://doi.org:10.1126/scitranslmed.aaf2968

90 Narla, R. K. et al. Modulation of CD47-SIRPα innate immune checkpoint axis with Fc-function detuned anti-CD47 therapeutic antibody. Cancer Immunology, Immunotherapy 71, 473–489 (2021). https://doi.org:doi:10.1007/s00262-021-03010-6

91 Jiang, Z., Sun, H., Yu, J., Tian, W. & Song, Y. Targeting CD47 for cancer immunotherapy. J Hematol Oncol 14, 180 (2021). https://doi.org:10.1186/s13045-021-01197-w

92 Dheilly, E. et al. Selective Blockade of the Ubiquitous Checkpoint Receptor CD47 Is Enabled by Dual-Targeting Bispecific Antibodies. Molecular Therapy 25, 523–533 (2017). https://doi.org:10.1016/J.YMTHE.2016.11.006

93 Trabulo, S., Aires, A., Aicher, A., Heeschen, C. & Cortajarena, A. L. Multifunctionalized iron oxide nanoparticles for selective targeting of pancreatic cancer cells. Biochimica et Biophysica Acta (BBA) - General Subjects 1861, 1597–1605 (2017). https://doi.org:10.1016/J.BBAGEN.2017.01.035

94 Chen, Q. et al. In situ sprayed bioresponsive immunotherapeutic gel for post-surgical cancer treatment. Nature Nanotechnology 2018 14:1 14, 89–97 (2018). https://doi.org:10.1038/s41565-018-0319-4

95 Chowdhury, S. et al. Programmable bacteria induce durable tumor regression and systemic antitumor immunity. Nature Medicine 2019 25:7 25, 1057–1063 (2019). https://doi.org:10.1038/s41591-019-0498-z

96 Li, Y. et al. A pH-dependent anti-CD47 antibody that selectively targets solid tumors and improves therapeutic efficacy and safety. Journal of Hematology & Oncology 2023 16:1 16, 1–21 (2023). https://doi.org:10.1186/S13045-023-01399-4

97 Xie, Y. J. et al. Improved antitumor efficacy of chimeric antigen receptor T cells that secrete single-domain antibody fragments. Cancer Immunology Research 8, 518–529 (2020). https://doi.org:10.1158/2326-6066.CIR-19-0734/470689/AM/IMPROVED-ANTI-TUMOR-EFFICACY-OF-CHIMERIC-ANTIGEN

98 La, H. T., Tran, D. B. T., Tran, H. M. & Nguyen, L. T. Third-Generation Anti-CD47-Specific CAR-T Cells Effectively Kill Cancer Cells and Reduce the Genes Expression in Lung Cancer Cell Metastasis. Journal of Immunology Research 2021 (2021). https://doi.org:10.1155/2021/5575260

99 Chen, H. et al. Delivery of CD47 blocker SIRPα-Fc by CAR-T cells enhances antitumor efficacy. Journal for ImmunoTherapy of Cancer 10, e003737–e003737 (2022). https://doi.org:10.1136/JITC-2021-003737

100 Dacek, M. M. et al. Potentiating antibody-dependent killing of cancers with CAR T cells secreting CD47-SIRPα checkpoint blocker. Blood (2023). https://doi.org:10.1182/BLOOD.2022016101

101 Chmielewski, M., Kopecky, C., Hombach, A. A. & Abken, H. IL-12 release by engineered T cells expressing chimeric antigen receptors can effectively Muster an antigen-independent macrophage response on tumor cells that have shut down tumor antigen expression. Cancer research 71, 5697–5706 (2011). https://doi.org:10.1158/0008-5472.CAN-11-0103

102 Pegram, H. J. et al. Tumor-targeted T cells modified to secrete IL-12 eradicate systemic tumors without need for prior conditioning. Blood 119 (2012). https://doi.org:10.1182/blood-2011-12-400044

103 Koneru, M., O’Cearbhaill, R., Pendharkar, S., Spriggs, D. R. & Brentjens, R. J. A phase I clinical trial of adoptive T cell therapy using IL-12 secreting MUC-16(ecto) directed chimeric antigen receptors for recurrent ovarian cancer. Journal of translational medicine 13 (2015). https://doi.org:10.1186/S12967-015-0460-X

104 Curran, K. J. et al. Enhancing antitumor efficacy of chimeric antigen receptor T cells through constitutive CD40L expression. Molecular therapy : the journal of the American Society of Gene Therapy 23 (2015). https://doi.org:10.1038/mt.2015.4

105 Kuhn, N. F. et al. CD40 Ligand-Modified Chimeric Antigen Receptor T Cells Enhance Antitumor Function by Eliciting an Endogenous Antitumor Response. Cancer cell 35 (2019). https://doi.org:10.1016/j.ccell.2019.02.006

106 Frigault, M. J. et al. Identification of chimeric antigen receptors that mediate constitutive or inducible proliferation of T cells. Cancer immunology research 3 (2015). https://doi.org:10.1158/2326-6066.CIR-14-0186

107 Chen, J. et al. NR4A transcription factors limit CAR T cell function in solid tumours. Nature 567 (2019). https://doi.org:10.1038/s41586-019-0985-x

108 Olaoba, O. T. et al. Is the new angel better than the old devil? Challenges and opportunities in CD47-SIRPα-based cancer therapy. Critical Reviews in Oncology/Hematology, 103939-103939 (2023). https://doi.org:10.1016/J.CRITREVONC.2023.103939

109 Deuse, T. et al. The SIRPα-CD47 immune checkpoint in NK cells. Journal of Experimental Medicine 218 (2021). https://doi.org:10.1084/JEM.20200839/211668

110 Labanieh, L., Majzner, R. G. & Mackall, C. L. Programming CAR-T cells to kill cancer. Nature Biomedical Engineering 2018 2:6 2, 377–391 (2018). https://doi.org:10.1038/s41551-018-0235-9

111 Lynn, R. C. et al. c-Jun overexpression in CAR T cells induces exhaustion resistance. Nature 2019 576:7786 576, 293–300 (2019). https://doi.org:10.1038/s41586-019-1805-z

112 Weber, E. W., Maus, M. V. & Mackall, C. L. Vol. 181 46-62 (Cell Press, 2020).

113 Weber, E. W. et al. Transient rest restores functionality in exhausted CAR-T cells through epigenetic remodeling. Science 372 (2021). https://doi.org:10.1126/SCIENCE.ABA1786/SUPPL_FILE/ABA1786_WEBER_SM.PDF

114 Freitas, K. A. et al. Enhanced T cell effector activity by targeting the Mediator kinase module. Science 378 (2022). https://doi.org:10.1126/SCIENCE.ABN5647/SUPPL_FILE/SCIENCE.ABN5647_MDAR_REPRODUCIBILITY_CHECKLIST.PDF

115 Komor, A. C., Kim, Y. B., Packer, M. S., Zuris, J. A. & Liu, D. R. Programmable editing of a target base in genomic DNA without double-stranded DNA cleavage. Nature 2015 533:7603 533, 420–424 (2016). https://doi.org:10.1038/nature17946

116 Gaudelli, N. M. et al. Programmable base editing of A•T to G•C in genomic DNA without DNA cleavage. Nature 2017 551:7681 551, 464–471 (2017). https://doi.org:10.1038/nature24644

117 Anzalone, A. V. et al. Search-and-replace genome editing without double-strand breaks or donor DNA. Nature 2019 576:7785 576, 149–157 (2019). https://doi.org:10.1038/s41586-019-1711-4

118 Hughes, M. S. et al. Transfer of a TCR Gene Derived from a Patient with a Marked Antitumor Response Conveys Highly Active T-Cell Effector Functions. https://home.liebertpub.com/hum 16, 457–472 (2005). https://doi.org:10.1089/HUM.2005.16.457

119 Ferrara, J., Guillen, F. J., Sleckman, B., Burakoff, S. J. & Murphy, G. F. Cutaneous acute graft-versus-host disease to minor histocompatibility antigens in a murine model: histologic analysis and correlation to clinical disease. The Journal of investigative dermatology 86, 371–375 (1986). https://doi.org:10.1111/1523-1747.EP12285612

120 Hao, Y. et al. Integrated analysis of multimodal single-cell data. Cell 184, 3573–3587.e3529 (2021). https://doi.org:10.1016/J.CELL.2021.04.048/ATTACHMENT/1E5EB5C1-59EE-4B2B-8BFA-14B48A54FF8F/MMC3.XLSX

121 Lim, S., Glasgow, J. E., Filsinger Interrante, M., Storm, E. M. & Cochran, J. R. Dual display of proteins on the yeast cell surface simplifies quantification of binding interactions and enzymatic bioconjugation reactions. Biotechnology Journal 12, 1600696–1600696 (2017). https://doi.org:10.1002/BIOT.201600696

122 Hunter, S. A. & Cochran, J. R. Vol. 580 21–44 (Academic Press Inc., 2016).

123 Hafemeister, C. & Satija, R. Normalization and variance stabilization of single-cell RNA-seq data using regularized negative binomial regression. Genome Biology 20, 1–15 (2019). https://doi.org:10.1186/S13059-019-1874-1/FIGURES/6

124 Chen, E. Y. et al. Enrichr: interactive and collaborative HTML5 gene list enrichment analysis tool. BMC bioinformatics 14 (2013). https://doi.org:10.1186/1471-2105-14-128

