## Extended Data for "Engineered CD47 protects T cells for enhanced antitumor immunity"

### Engineered CD47 protects T cells for enhanced antitumor immunity: Extended Data Figures

#### Extended Data Figure 1

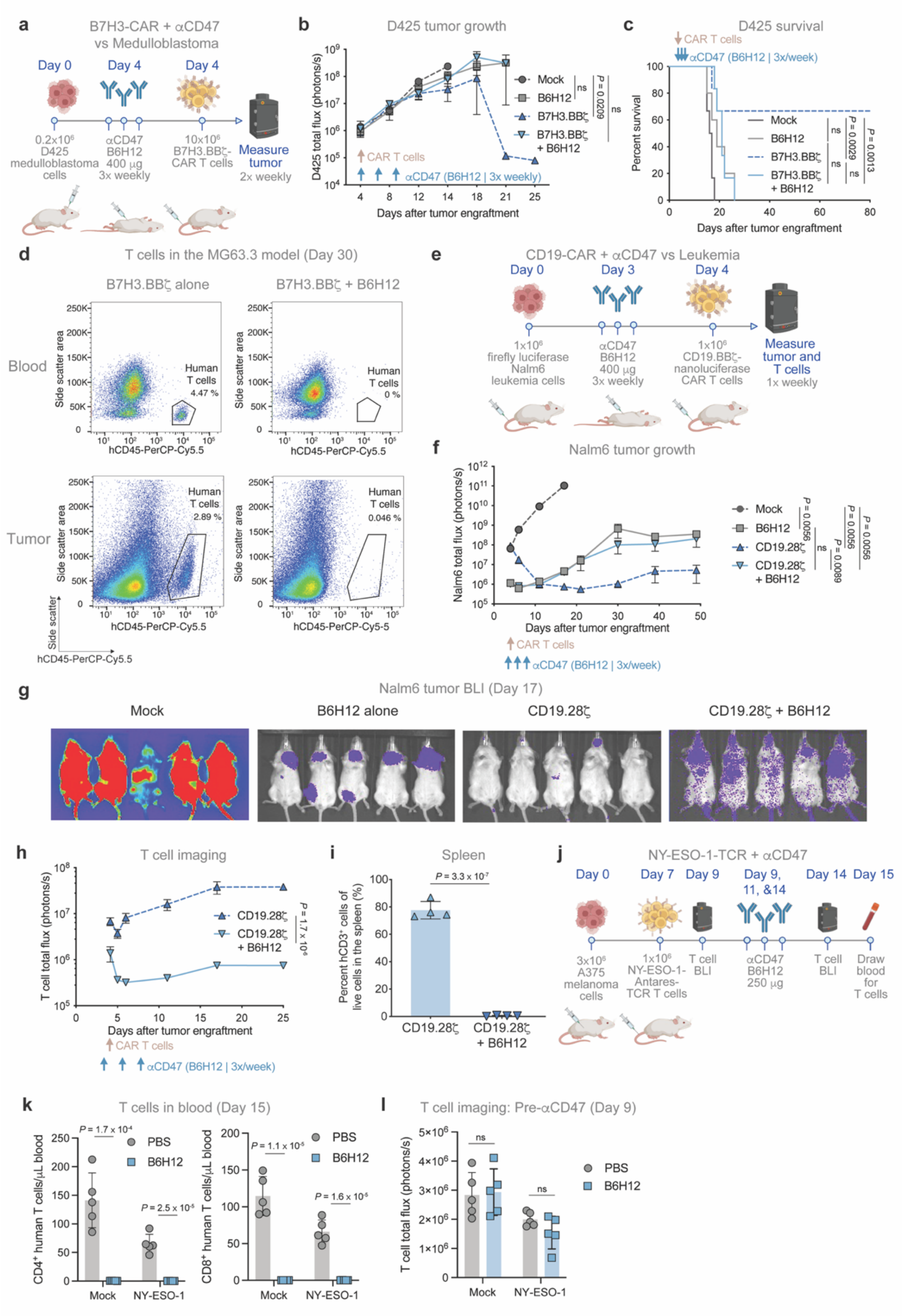

**Extended Data Figure 1: αCD47 therapy blunts CAR T efficacy by depleting CAR T cells**, related to Figure 1

(a – c) D425 medulloblastoma model.

(a) D425 treatment scheme. Mice engrafted with $0.2\times10^6$ D425 cells in the cerebellum were treated ± B6H12 intraperitoneally (IP) three times per week (400 µg/dose) starting on day 4. Mice were also treated intravenously (IV) with $10\times10^6$ mock or B7H3.BBζ-CAR T cells on day 4.

(b) Quantification of D425 tumor growth by BLI, treated as described in (a). Data are the mean ± SEM of n = 5 mice/arm. Data are representative of two independent experiments.

(c) Survival of mice in the D425 model shown in (b). n = 5 mice per treatment arm.

(d) Representative flow cytometry plots of $hCD45^+$ T cells identified in the blood and tumor in the MG63.3 model on day 30 post tumor engraftment, treated with $10\times10^6$ B7H3.BBζ-CAR T cells on day 15 ± 3 doses of B6H12 treatment (400 µg/dose).

(e – i) Nalm6 leukemia model.

(e) Nalm6 model treatment scheme. Mice engrafted IV with $1\times10^6$ Nalm6-fLuc cells were treated ± B6H12 (400 µg/dose; IP) three times per week starting on day 3. Mice were then treated IV with $1\times10^6$ mock or CD19.28ζ-CAR T cells on day 4.

(f) Quantification of Nalm6 tumor growth by BLI, treated as described in (e). Data are the mean ± SEM of n = 5 mice/arm.

(g) Nalm6 tumor BLI on day 17, treated as described in (e).

(h) Quantification of T cell BLI in the Nalm6 model, treated as described in (e). Data are the mean ± SEM of n = 5 mice/arm.

(i) Quantification of T cells by flow cytometry from the spleen in the Nalm6 model, treated as described in (e). Data are the mean ± SD of n = 5 mice.

(j) A375 – NY-ESO-1 treatment scheme. Mice engrafted subcutaneously (SQ) with $3\times10^6$ A375 were treated with $1\times10^6$ mock-Antares or NY-ESO-1-Antares-TCR T cells IV on day 7 ± three doses of B6H12 (250 µg/dose; IP) on days 9, 11, and 14. Mice were imaged by BLI before (day 9) and after (day 14) αCD47 treatment. Blood was collected on day 15.

(k) Quantification of $hCD4^+$ (left) and $hCD8^+$ (right) T cells by flow cytometry from the blood on day 15 in the A375 – NY-ESO-1 model, treated as described in (j). Data are the mean ± SD of n = 5 mice.

(l) Quantification of T cell BLI in the A375 – NY-ESO-1 model prior to B6H12 treatment on day 9, treated as described in (j). Mock is a shared control duplicated in T cell depletion in Extended Data Figure 7a. Data are the mean ± SD of n = 5 mice.

[(b) and (h)] Two-way analysis of variance (ANOVA) test with Tukey's multiple comparison test. ns = not significant.
[(c)] Log-rank Mantel-Cox test. ns = not significant.
[(f – day 17), (i), (k), and (l)] Unpaired two-tailed Student's *t* test. ns = not significant.

#### Extended Data Figure 2

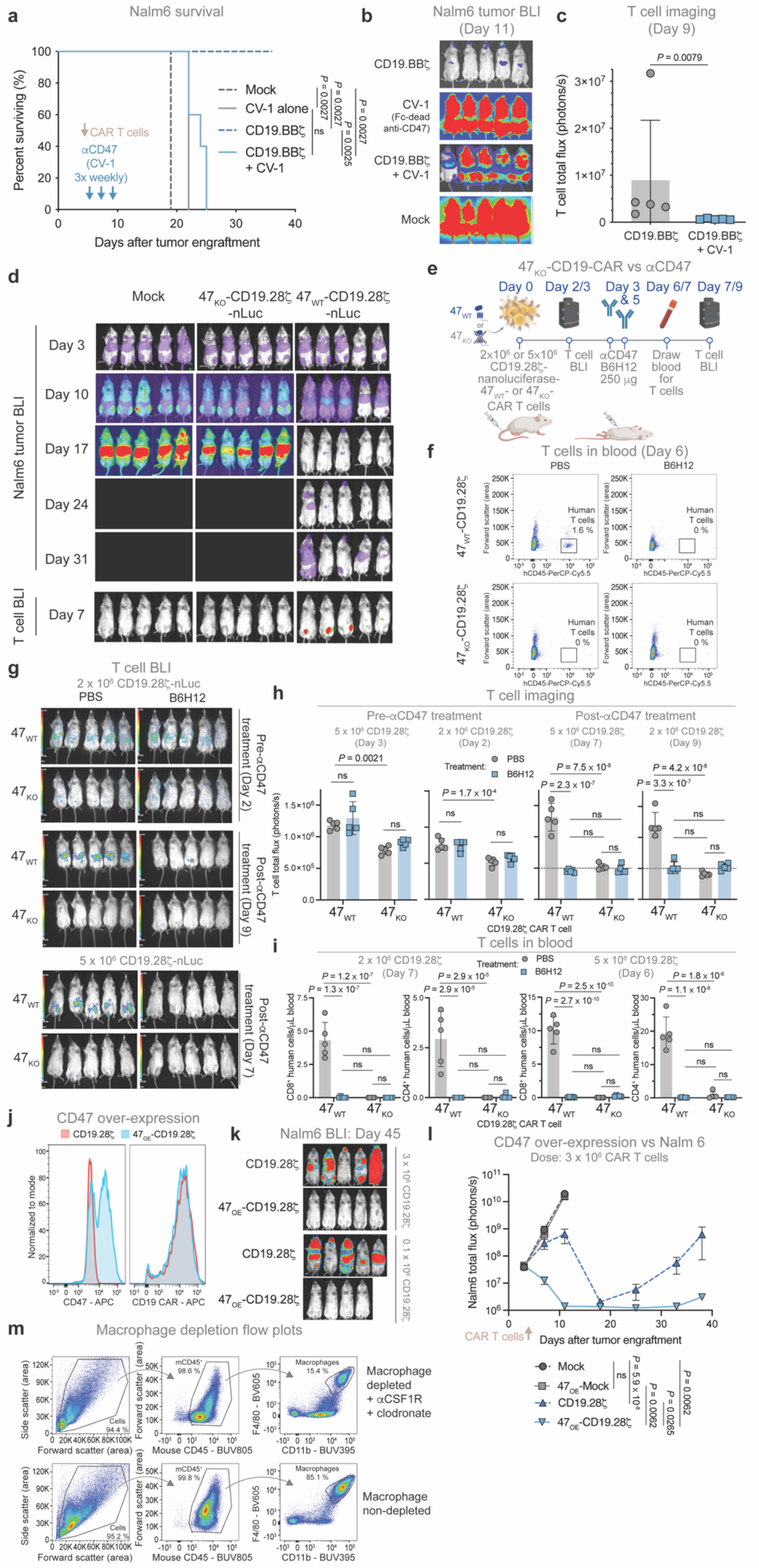

**Extended Data Figure 2: CAR T depletion after $\alpha$CD47 treatment phenocopies CD47 KO and is mediated by macrophages**, related to Figures 1 & 2

(a – c) Nalm6 leukemia model treated with CAR T cells and CV-1.

(a) Survival of mice in the Nalm6 – CV-1 model shown in Fig. 1g. n = 5 mice per treatment arm.

(b) Nalm6 BLI on day 11 in the Nalm6 – CV-1 model, treated as described in Fig. 1f.

(c) Quantification of T cell BLI on day 9 after CV-1 treatment in the Nalm6 model, treated as described in Fig. 1f.

(d) Nalm6 tumor progression (top) and T cells on day 7 (bottom) by BLI in the Nalm6 – $47_{KO}$-CAR T model, treated as described in Fig. 2b.

(e – i) T cell depletion by CD47 KO ($47_{KO}$) or B6H12 treatment.

(e) T cell depletion model scheme. Non-tumor bearing mice treated IV with $5\times10^6$ or $2\times10^6$ $47_{KO}$-CD19.28$\zeta$-nLuc-CAR T cells in separate experiments, with ($47_{WT}$) or without ($47_{KO}$) CD47 exogenous expression, were then treated twice ± B6H12 (250 µg/dose; IP) on days 3 and 5. Mice were imaged by BLI before (day 2 or 3) and after (day 7 or 9) $\alpha$CD47 treatment, and had blood drawn on day 6 or 7.

(f) Example flow cytometry plots of $hCD45^+$ T cells identified in the blood in the T cell depletion model, treated as described in (e).

(g) T cell BLI in the T cell depletion model described in (e), following IV treatment with $2\times10^6$ (top) or $5\times10^6$ (bottom) $47_{WT}$- or $47_{KO}$-CD19.28$\zeta$-nLuc-CAR T cells. $47_{WT}$ is a shared condition duplicated in T cell depletion in Extended Data Fig. 5e.

(h) Quantification of T cell BLI in the T cell depletion model, treated as described in (e), with mice treated IV with $2\times10^6$ or $5\times10^6$ $47_{WT}$- or $47_{KO}$-CD19.28$\zeta$-nLuc-CAR T cells, as indicated. Dashed line indicates limit of detection. Data are the mean ± SD of n = 5 mice. $47_{WT}$ is a shared condition duplicated in T cell depletion in Extended Data Fig. 5f.

(i) Quantification of $CD8^+$ (left) and $CD4^+$ (right) T cells in the blood on day 6 or 7 by flow cytometry in the T cell depletion model, treated as described in (e), with mice treated IV with $2\times10^6$ or $5\times10^6$ $47_{WT}$- or $47_{KO}$-CD19.28$\zeta$-nLuc-CAR T cells in separate experiments. Data are the mean ± SD of n = 5 mice. $47_{WT}$ is a shared condition duplicated in T cell depletion in Extended Data Figure 5g.

(j) CD47 (left) and CAR (right) expression on T cells by flow cytometry after CD47 OE ($47_{OE}$). Data are representative of n > 3 donors.

(k) Nalm6 BLI on day 45 after dosing with $3\times10^6$ (top) or $0.1\times10^6$ (bottom) CAR T cells. Mice engrafted IV with $1\times10^6$ Nalm6 were treated IV with $3\times10^6$ or $0.1\times10^6$ mock, $47_{OE}$-mock, CD19.28$\zeta$-, or $47_{OE}$-CD19.28$\zeta$-CAR T cells on day 4 in separate experiments with different CAR T doses.

(l) Quantification of Nalm6 tumor growth by BLI. Mice engrafted IV with $1\times10^6$ Nalm6 were treated IV with $3\times10^6$ mock, $47_{OE}$-mock, CD19.28$\zeta$-, $47_{OE}$-CD19.28$\zeta$-CAR T cells on day 4. Data are the mean ± SEM of n = 5 mice/arm. Mock and $47_{OE}$-mock are a shared conditions duplicated in Fig. 2e.

(m) Representative flow plots and gating strategy for detection of macrophages in samples collected following peritoneal lavage.

[(a)] Log-rank Mantel-Cox test. **$P$ < 0.001.
[(c)] Mann-Whitney test. **$P$ < 0.01.
[(h), (i), and (l)] Two-way analysis of variance (ANOVA) test with Tukey's multiple comparison test. ns = not significant.

### Extended Data Figure 3

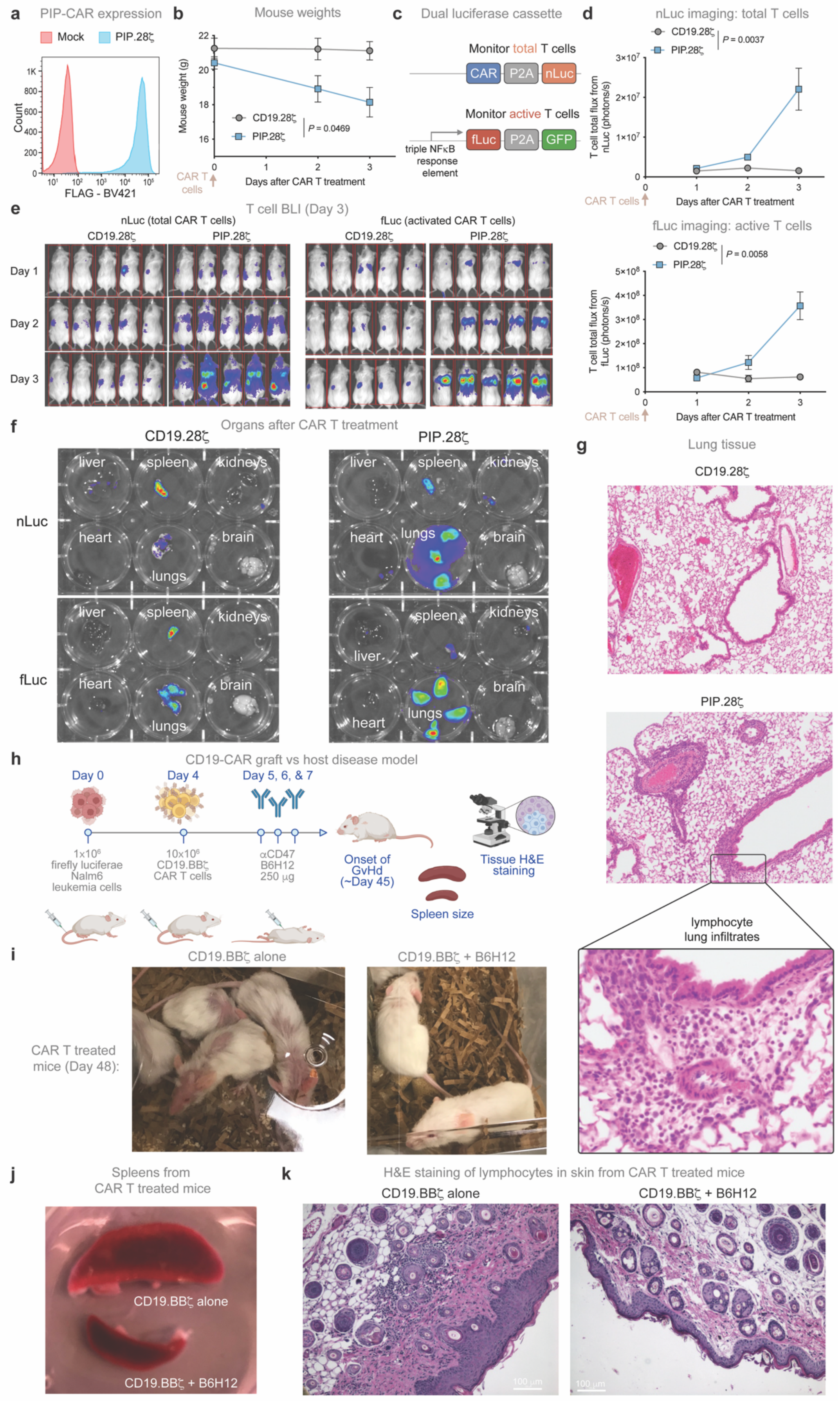

**Extended Data Figure 3: αCD47 can limit toxicities from a pan-tumor integrin targeting PIP-CAR and GvHD**, related to Figure 4

(a – g) PIP-CAR toxicity model

(a) PIP-CAR expression with CD28 costimulatory domain by flow cytometry. Data are representative of n = 3 donors.

(b) Weights of mice following treatment with $5\times10^6$ CD19.28ζ- or PIP.28ζ-CAR T cells IV. Data are the mean ± SEM of n = 5 mice/arm.

(c) Cartoon of the dual luciferase reporter design for tracking CAR T activity *in vivo*. nLuc is linked to CAR expression, fLuc is linked to NFκB activation.

(d) Quantification of (top) nLuc and (bottom) fLuc T cell BLI, treated as described in (b). Data are the mean ± SEM of n = 5 mice/arm.

(e) BLI detecting total T cells (nLuc, left) and activated T cells (fLuc, right) following IV treatment with $5\times10^6$ CD19.28ζ- or PIP.28ζ-CAR T cells 3 days after infusion.

(f) Representative images of nLuc (top) and fLuc (bottom) BLI from organs extracted from mice treated as described in (e), four days after CAR T administration.

(g) Representative hematoxylin and eosin (H&E) stained images of lung tissue collected from mice treated as described in (e) four days after CAR T treatment. Bottom panel is an expanded view of the boxed region in the middle panel.

(h – k) CAR T cell-induced GvHD model

(h) Scheme of GvHD model. Mice engrafted IV with $1\times10^6$ Nalm6-fLuc were treated IV with a high dose of $10\times10^6$ CD19.BBζ-CAR T cells on day 4 ± three doses of B6H12 (250 µg/dose; IP) on days 5, 6, and 7. Mice were monitored for onset of GvHD around day 45.

(i – k) Representative images of (i) mice demonstrating GvHD derived alopecia (day 48 post-CAR T infusion), (j) spleens from mice (collected on day 48 post-CAR T infusion), and (k) H&E staining of skin sections (collected on day 48 post-CAR T infusion) from mice treated as described in (h).

[(b) and (d)] Two-way analysis of variance (ANOVA) test with Tukey's multiple comparison test. ns = not significant.

### Extended Data Figure 4

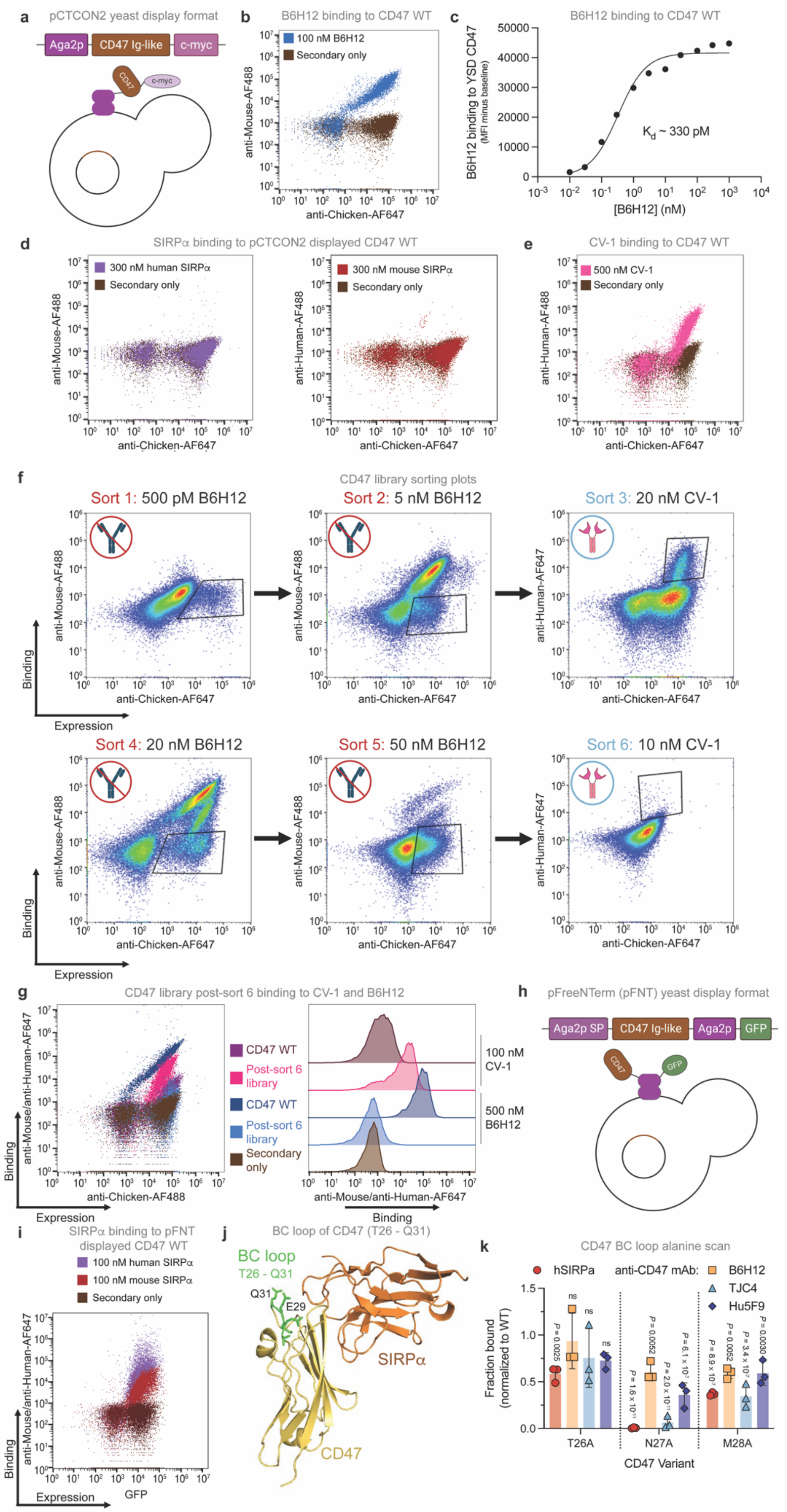

**Extended Data Figure 4: Engineered variants of CD47 retain SIRPα binding and demonstrate a loss of binding to some, but not all, αCD47 antibodies**, related to Figure 5

(a) Cartoon of yeast displayed CD47 Ig-like domain using the pCTCON2 vector. CD47 is displayed as an N-terminal fusion.

(b) Binding of 100 nM B6H12 to yeast displayed CD47 in pCTCON2 by flow cytometry. Data are representative of n = 3 independent experiments.

(c) Binding curve of B6H12 to yeast displayed CD47 in pCTCON2, measured over multiple concentrations by flow cytometry. Data are the MFI of n = 1 experiment.

(d) Binding of 300 nM human (left) and mouse (right) SIRPα to yeast displayed CD47 in pCTCON2 by flow cytometry. Data are representative of n = 3 independent experiments.

(e) Binding of 500 nM CV-1 to yeast displayed CD47 in pCTCON2 by flow cytometry. Data are representative of n = 3 independent experiments.

(f) Flow cytometry sorting plots of all six sorts of the CD47 library, indicating negative sorts to B6H12 and positive sorts to CV-1. Collected population indicated by the black box in each plot.

(g) Binding of 500 nM B6H12 or 100 nM CV-1 to the yeast displayed CD47 library population collected after sort 6 or yeast displayed WT CD47. Data are representative of n = 2 independent experiments.

(h) Cartoon of yeast-displayed CD47 Ig-like domain using the pFreeNTerm (pFNT) vector. CD47 is displayed as a C-terminal fusion, along with GFP to monitor protein expression.

(i) Binding of 100 nM human and mouse SIRPα to yeast displayed CD47 in pFNT by flow cytometry. Data are representative of n = 3 independent experiments.

(j) Crystal structure of CD47 (yellow) binding SIRPα (orange) [PDB: 2JJS], identifying the CD47 BC loop (green), containing CD47 residues T26 – Q31.

(k) Binding of 100 nM B6H12, 100 nM TJC4, 100 nM Hu5F9, and 100 nM hSIRPα to yeast displayed $47_{T26A}$, $47_{N27A}$, and $47_{M28A}$ variants. Data are the mean ± SD of n = 3 individual yeast clones, normalized to MFI from binding to $47_{WT}$.

[(k)] Two-way analysis of variance (ANOVA) test with Tukey's multiple comparison test. ns = not significant. Comparison is between indicated group and binding to CD47 WT expressing cells.

**Extended Data Figure 5**

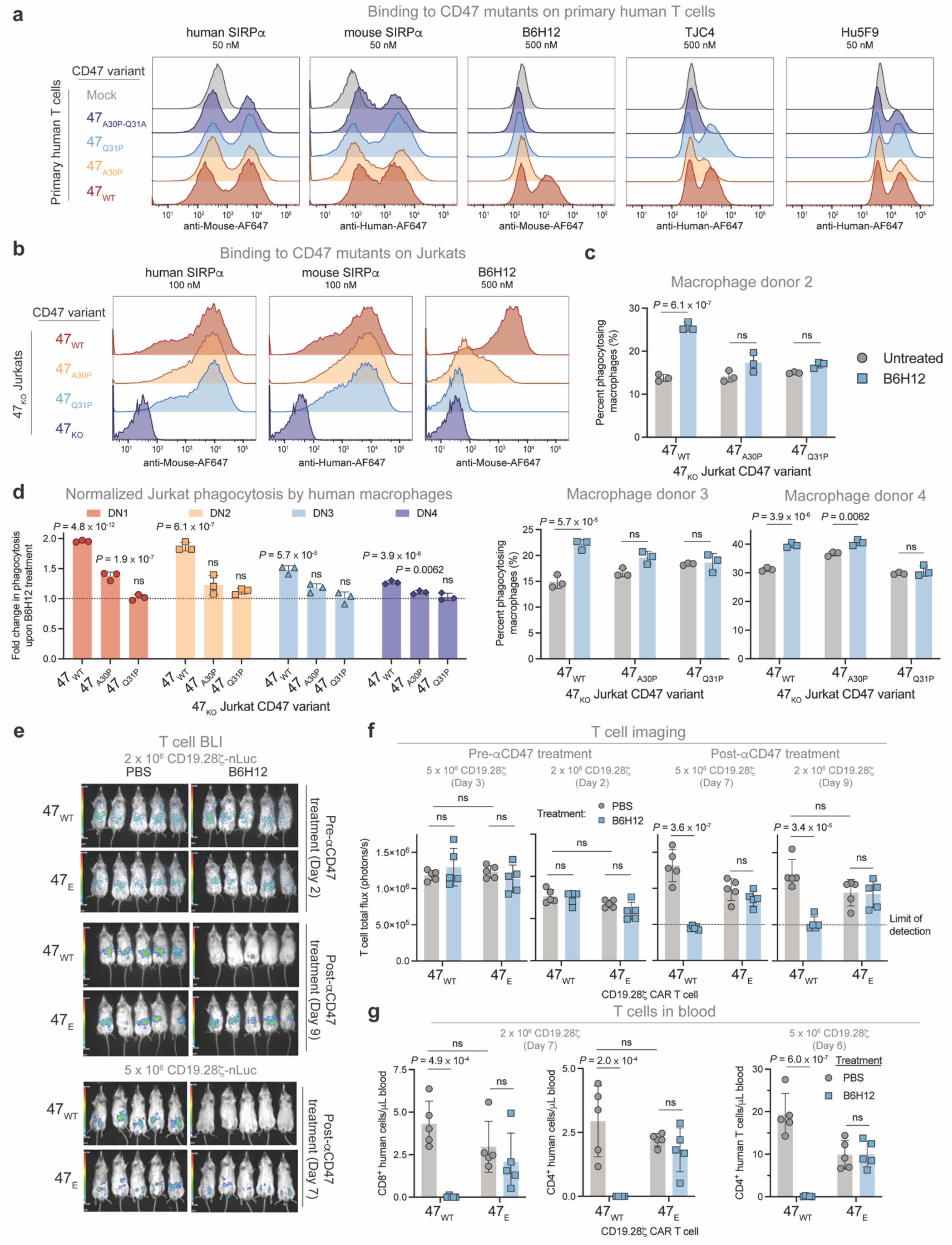

**Extended Data Figure 5: Expression of $47_E$ on T cells mitigates B6H12 induced phagocytosis due to lack of antibody binding *in vitro* and *in vivo***, related to Figure 5

(a) Representative flow cytometry histograms of 50 nM hSIRPα, 50 nM mSIRPα, 500 nM B6H12, 500 nM TJC4, and 50 nM Hu5F9 binding to $47_{WT}$, $47_{A30P}$, $47_{Q31P}$, and $47_{A30P\text{-}Q31A}$ over-expressed on primary human T cells. Data are representative of n = 3 independent experiments.

(b) Representative flow cytometry histograms of 100 nM hSIRPα, 100 nM mSIRPα, and 500 nM B6H12 binding to $47_{WT}$, $47_{A30P}$, and $47_{Q31P}$ expressed on Jurkats with endogenous $47_{KO}$. Data are representative of n = 3 independent experiments.

(c and d) Quantification of phagocytosis by primary human macrophages from multiple donors of CSFE labeled Jurkats with endogenous $47_{KO}$, expressing $47_{WT}$, $47_{A30P}$, or $47_{Q31P}$ variants after one hour of co-culture. Data are the mean ± SD of triplicate wells (n = 3).

(e) T cell BLI in the $47_E$-T cell depletion model described in Fig. 5k, following IV treatment with $2\times10^6$ (top) or $5\times10^6$ (bottom) $47_{WT}$- or $47_E$-CD19.28ζ-nLuc-CAR T cells. $47_{WT}$ is a shared condition duplicated in T cell depletion in Extended Data Fig. 2g.

(f) Quantification of T cell BLI in the $47_E$-T cell depletion model before and after αCD47 treatment, treated as described in Fig. 5k, with mice treated IV with $2\times10^6$ or $5\times10^6$ $47_{WT}$- or $47_E$-CD19.28ζ-nLuc-CAR T cells, as indicated. Data are the mean ± SD of n = 5 mice. $47_{WT}$ is a shared condition duplicated in T cell depletion in Extended Data Fig. 2h.

(g) Quantification of $CD8^+$ (left) and $CD4^+$ (right) T cells in the blood on day 6 by flow cytometry in the $47_E$-T cell depletion model, treated as described in Fig. 5k, with mice treated with $2\times10^6$ or $5\times10^6$ $47_{WT}$- or $47_E$-CD19.28ζ-nLuc-CAR T cells, as indicated. Data are the mean ± SD of n = 5 mice. $47_{WT}$ is a shared condition duplicated in T cell depletion in Extended Data Fig. 2i.

[(c), (d), (f), and (g)] Two-way analysis of variance (ANOVA) test with Tukey's multiple comparison test. ns = not significant. Significance in (d) indicates the comparison between each condition ± B6H12.

#### Extended Data Figure 6

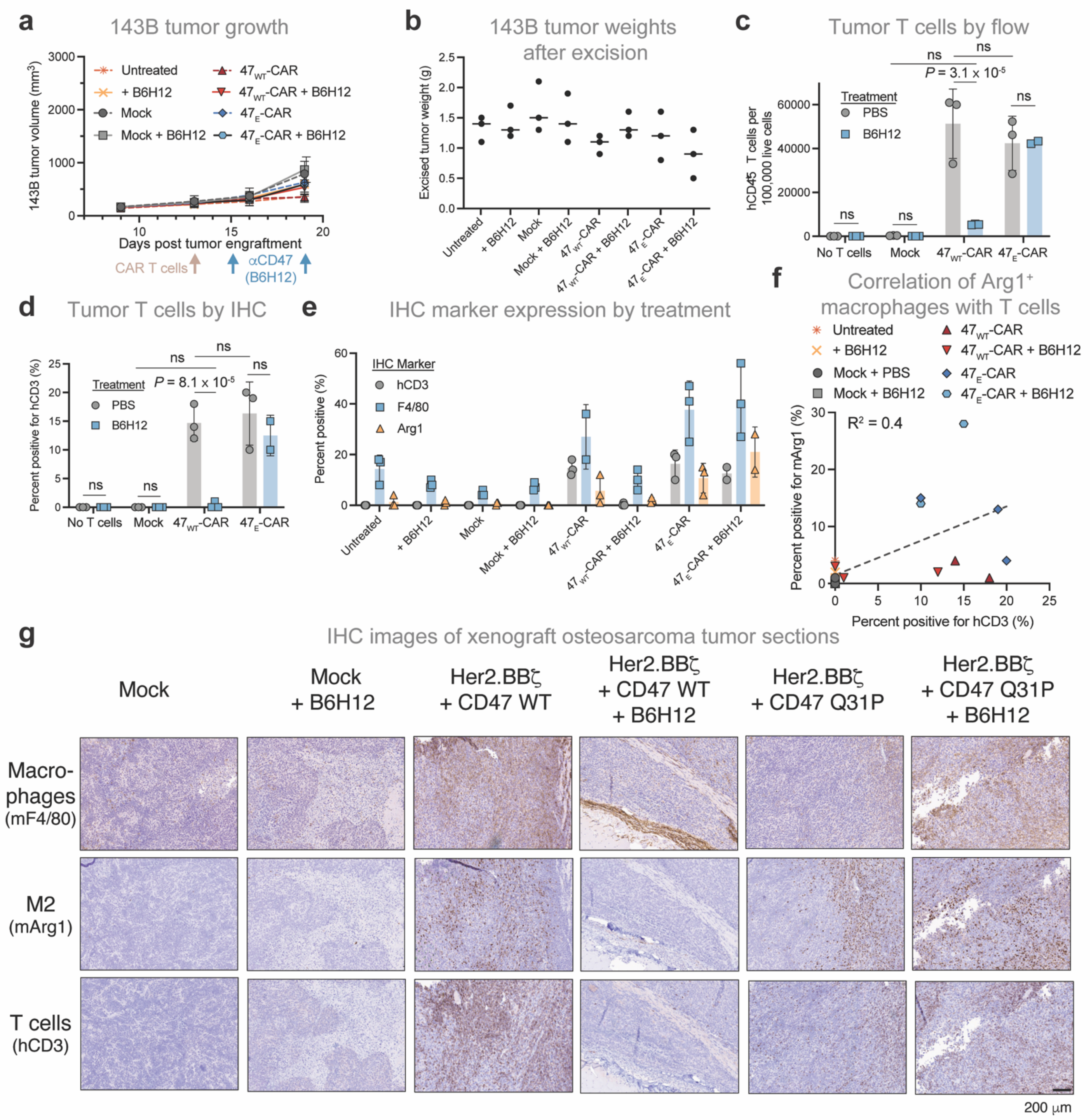

**Extended Data Figure 6: CAR T treatment increases macrophage tumor infiltration**, related to Figure 6

(a) 143B tumor growth over treatment period for correlative study, treated as described in Fig. 6a. Data are the mean ± SD of n = 3 mice/arm.

(b) 143B excised tumor weights after 8 days of treatment, treated as described in Fig. 6a. Data are the mean of n = 3 mice.

(c) Human $CD45^+$ T cells identified by flow cytometry of dissociated tumors, treated as described in Fig. 6a. Data are the mean ± SD of n = 2 ($47_E$-CAR + B6H12) or n = 3 (all other samples) mice.

(d) Quantification of hCD3 staining of IHC sections produced from tumor sections, treated as described in Fig. 6a. Data are the mean ± SD of n = 3 mice.

(e) Quantification of human CD3 (T cells), mouse Arg1 (M2 macrophages), and mouse F4/80 (total macrophages) staining of IHC sections produced from tumor sections, treated as described in Fig. 6a. Data are the mean ± SD of n = 2 – 3 mice.

(f) Correlation of quantification of hCD3 and mArg1 staining in IHC sections of tumors treated as described in Fig. 6a. Data points are representative of individual tumors, colored by treatment group (n = 23). $R^2$ calculated by simple linear regression.

(g) Representative images of IHC tumor sections stained for mF4/80, mArg1, and hCD3, treated as described in Fig. 6a.

[(c) and (d)] Two-way ANOVA test with Tukey's multiple comparison test. ns = not significant.

### Extended Data Figure 7

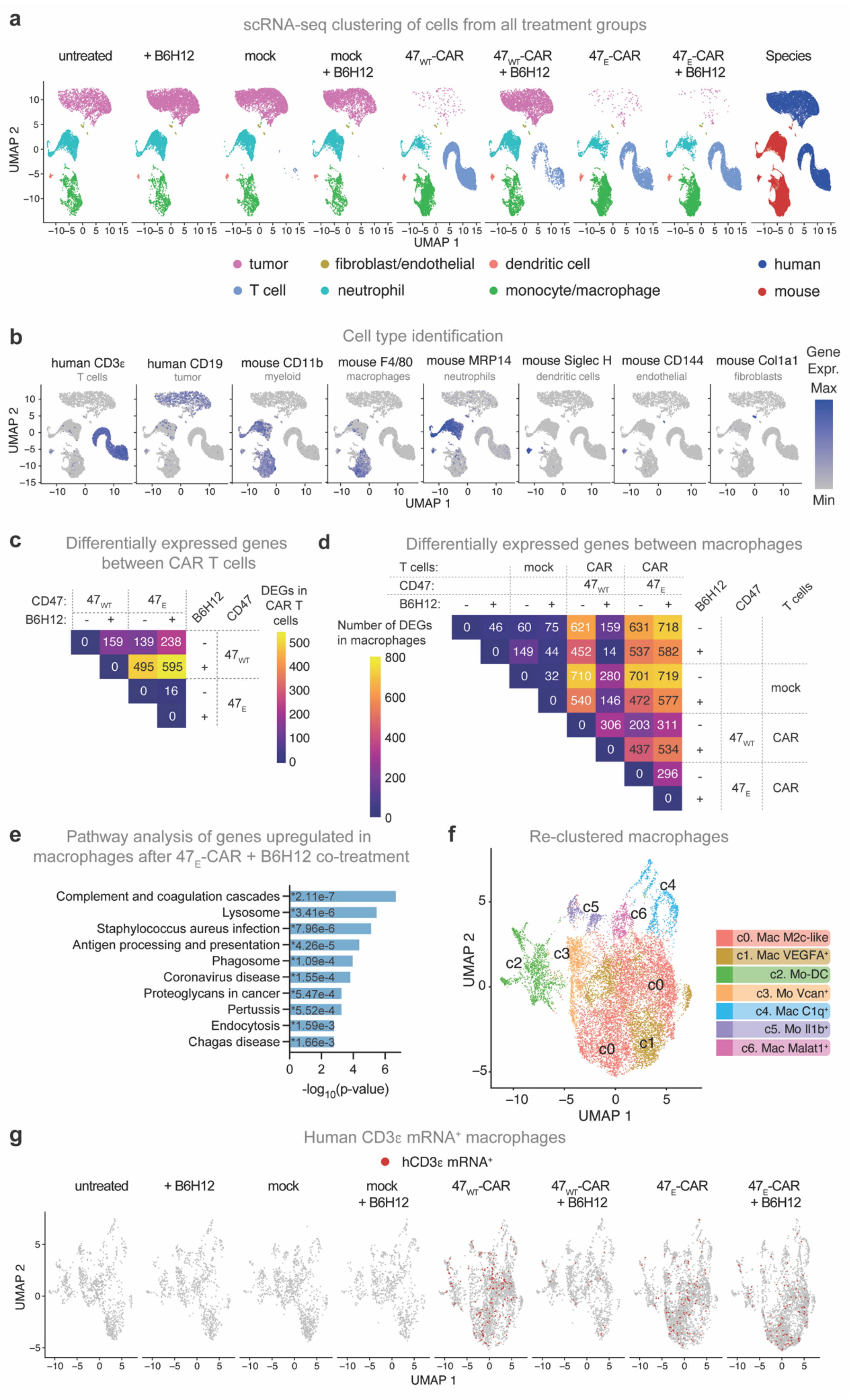

**Extended Data Figure 7: CAR T treatment results in unique populations of tumor infiltrating macrophages**, related to Figure 6

(a and b) scRNA-seq profile of dissociated tumor and infiltrating immune cells. Dots represent individual cells. *n* = 53,062 cells from 8 experimental conditions with three mice per treatment group, colored by (a) cell type (left eight plots; UMAPs represent distinct treatment conditions), species (far right), or (b) gene expression level.

(c) Comparison of differentially expressed genes between CAR T cells of different treatment groups described in Fig. 6a. Statistical significance was determined with Seurat; **P* adj < 0.05.

(d) Comparison of differentially expressed genes between macrophages of different treatment groups described in (a). Statistical significance was determined with Seurat; **P* adj < 0.05.

(e) Enrichr pathway analysis of the top 100 upregulated genes in tumor infiltrating macrophages [monocyte/macrophage cluster in (a)] in $47_E$-CAR + B6H12 treated tumors compared with untreated controls. The KEGG Human Pathway collection was queried with converted murine gene IDs.

(f) UMAP of the identified macrophage/monocyte population in (a), subsetted and re-clustered, colored by cluster. Dots represent individual cells. *n* = 13,082 cells from 8 experimental conditions.

(g) UMAP of the re-clustered macrophage/monocyte population from (f), colored for red if hCD3ε mRNA expression > 0. Dots represent individual cells. *n* = 13,082 cells from 8 experimental conditions, with n = 350 total cells identified as hCD3ε $mRNA^+$. UMAPs represent distinct treatment conditions.

See also Supplementary Tables 1 and 2.

#### Extended Data Figure 8

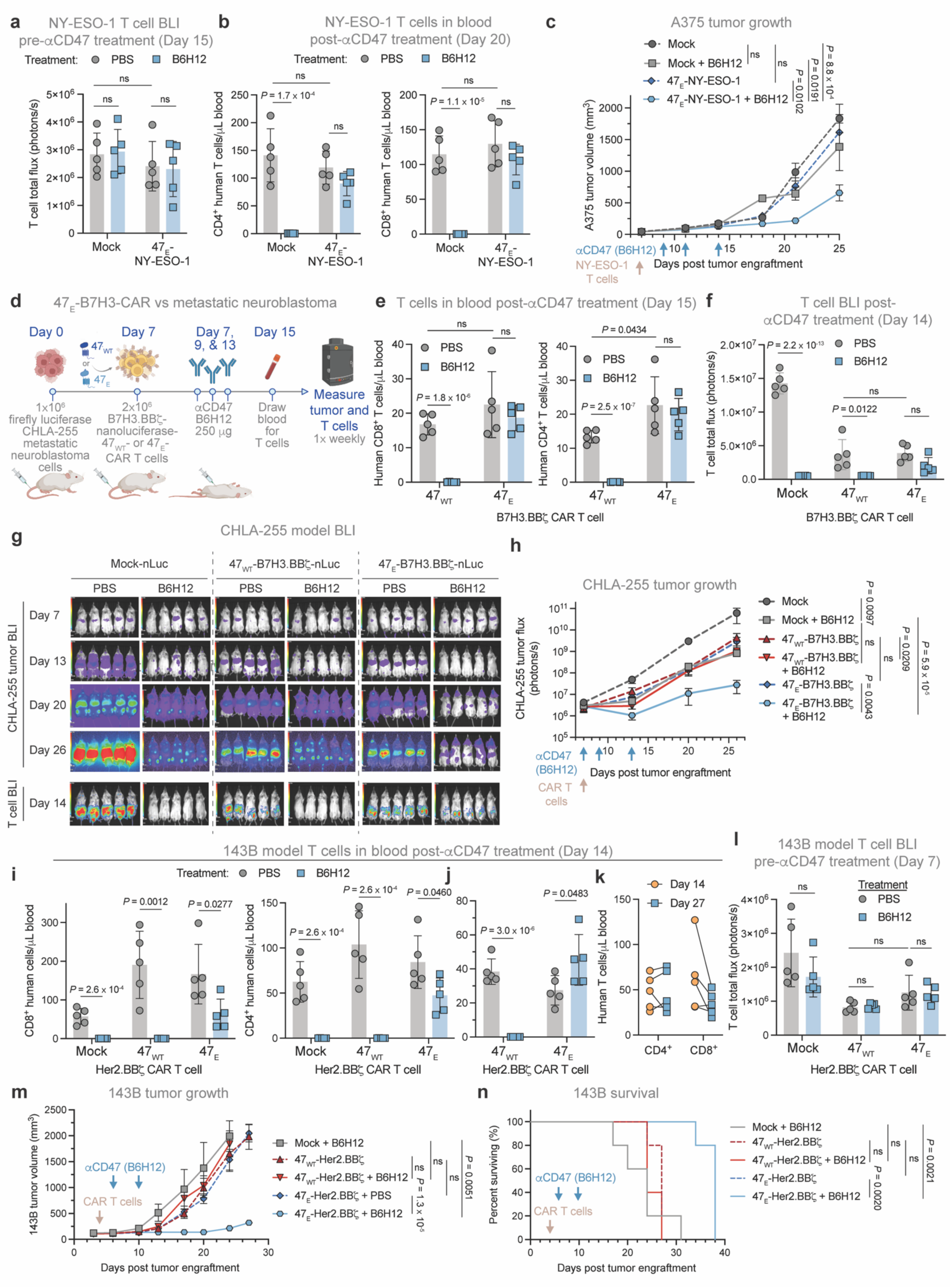

**Extended Data Figure 8: Expression of $47_E$ on tumor-targeting T cells permits pairing with αCD47 therapy and results in improved tumor control**, related to Figure 7

(a) Quantification of T cells by BLI in the A375 – NY-ESO-1 model before αCD47 treatment (day 9). Mice engrafted SQ with $3\times10^6$ A375 were treated IV with $1\times10^6$ mock-Antares or $47_E$-NY-ESO-1-Antares-TCR T cells (with endogenous $47_{KO}$) on day 7 ± three doses of B6H12 (250 µg/dose; IP) on days 9, 11, and 14. Mice were imaged by BLI before (day 9) and after (day 14) αCD47 treatment. Blood was collected on day 15. Data are the mean ± SD of n = 5 mice. Mock is a shared control duplicated in T cell depletion in Extended Data Figure 1l.

(b) Quantification of $hCD4^+$ (left) and $hCD8^+$ (right) T cells by flow cytometry from the blood in the A375 – $47_E$-NY-ESO-1 model treated as described in (a). Data are the mean ± SD of n = 5 mice. Mock is a shared control duplicated in T cell depletion in Extended Data Figure 1k.

(c) A375 tumor growth treated as described in (a), with T cells derived from a different donor than shown in Fig. 7d. Data are the mean ± SEM of n = 5 mice/arm.

(d) Schematic of CHLA-255 metastatic neuroblastoma model treatment. Mice were engrafted IV with $1\times10^6$ CHLA-255-fLuc cells and treated IV with $2\times10^6$ mock-nLuc, $47_{WT}$- or $47_E$-B7H3.BBζ-nLuc-CAR T cells (with endogenous $47_{KO}$) on day 7. Mice were then treated ± three doses of B6H12 (250 µg/dose; IP) on days 7, 9 and 13. T cells were imaged by BLI on day 14 and blood was collected on day 15.

(e) Quantification of $hCD8^+$ (left) and $hCD4^+$ (right) T cells by flow cytometry from the blood on day 15 in the CHLA-255 model, treated as described in (d). Data are the mean ± SD of n = 5 mice.

(f) Quantification of T cell BLI on day 14 in the CHLA-255 model, treated as described in (d). Data are the mean ± SD of n = 5 mice.

(g) CHLA-255 tumor progression (top) and T cells on day 14 (bottom) by BLI in the CHLA-255 model, treated as described in (d).

(h) Quantification of CHLA-255 tumor growth by BLI, treated as described in (d). Data are the mean ± SEM of n = 5 mice/arm.

(i and j) Quantification of $CD8^+$ (left, j) and $CD4^+$ (right, j & k) human T cells derived from two independent donors (represented by panels j & k) in the blood on day 14 by flow cytometry after B6H12 treatment in the 143B model, treated as described in Fig. 7e. Data are the mean ± SD of n = 5 mice.

(k) Quantification of human $CD4^+$ and $CD8^+$ T cells in the blood on day 14 and day 27 of tumor growth in the 143B model, treated as described in Fig. 7e. Data represent the values of n = 5 mice.

(l) Quantification of T cell BLI prior to B6H12 treatment in the 143B model, treated as described in Fig. 7e. Data are the mean ± SD of n = 5 mice.

(m and n) 143B tumor (m) growth and (n) survival in the 143B model, treated as described in Fig. 7e, using T cells derived from a different donor than Fig. 7i. Data are (m) the mean ± SEM or (n) representative of n = 5 mice.

[(a), (b), (c), (e), (f), (h), (i), (j), (l), and (m)] Two-way ANOVA test with Tukey's multiple comparison test. ns = not significant.
[(n)] Log-rank Mantel-Cox test. ns = not significant.
